# PAF1 and FACT cooperate with MLL-AF4 to drive enhancer activity in leukemia

**DOI:** 10.1101/2022.05.27.493676

**Authors:** Nicholas T. Crump, Alastair Smith, Laura Godfrey, Nicole Jackson, Siobhan Rice, Jaehoon Kim, Venkatesha Basrur, Damian Fermin, Kojo Elenitoba-Johnson, Robert G. Roeder, C. David Allis, Irene Roberts, Anindita Roy, Thomas A. Milne

## Abstract

Aberrant enhancer activation has been identified as a key mechanism driving oncogene expression in many cancers. Here we use TOPmentation (Transcription factor-OPtimized ChIPmentation) to probe enhancer usage in primary MLL-rearranged acute lymphoblastic leukemia. We find that MLL-AF4, commonly held to promote transcription by binding to gene promoters, is also present at many active enhancers, where it assembles a complex of transcriptional co-activators normally found in the gene body. This includes DOT1L, ENL, PAF1, and a newly identified interaction with the histone chaperone FACT. By chemical degradation, we demonstrate that PAF1 and FACT are required for enhancer activity, including maintaining histone H3K27 acetylation, enhancer RNA transcription and enhancer-promoter interactions. This work identifies novel roles for PAF1 and FACT in enhancer function, and reveals an enhancer-targeting mechanism by which MLL-AF4 upregulates transcription, recruiting transcription machinery through a network of multivalent interactions to control enhancer activity and gene expression in acute leukemias.

## Introduction

Enhancers are key regulatory elements that contribute to gene expression. Aberrant enhancer activity is a major factor in many cancers, often involving specific DNA mutations or even large-scale DNA rearrangements (Barwick et al., 2019; Mansour et al., 2014). Epigenetic changes in cancer are also increasingly recognized as important drivers of gene expression changes in a mutation-independent manner (Hanahan, 2022). Work has mainly focused on activity at promoters, but epigenetic changes can also be a driving force in enhancer dysregulation without mutations in the underlying sequence (Bradner et al., 2017). Many enhancer-associated factors, for example p300/CBP, are drivers of oncogenesis (Attar and Kurdistani, 2017; Farria et al., 2015; Lasko et al., 2017), and transcription factors (TFs) that bind at enhancers are often upregulated in cancers (Bradner et al., 2017).

Rearrangements of the *Mixed Lineage Leukemia* (*MLL*/*KMT2A*) gene (*MLL*r) cause aggressive, poor prognosis acute lymphoblastic (ALL) and acute myeloid (AML) leukemias in both children and adults (Krivtsov and Armstrong, 2007; Milne, 2017; Rice and Roy, 2020) and are associated with very few cooperating mutations (Agraz-Doblas et al., 2019; Andersson et al., 2015; Bill et al., 2020; Cancer Genome Atlas Research et al., 2013; Dobbins et al., 2013). Despite the relatively simple genetic landscape, *MLL*r leukemias exhibit a large number of epigenetic and transcriptional changes, suggesting that the MLL fusion protein (MLL-FP) drives oncogenesis via transcriptional reprogramming, and therefore provides a good model to study aberrant activation of enhancers.

Active enhancers commonly display characteristics including an open chromatin conformation and post-translational histone modifications such as histone H3 lysine-4 monomethylation (H3K4me1) and H3 lysine-27 acetylation (H3K27ac) (Rada-Iglesias et al., 2011). Exactly how enhancers control gene expression remains unknown, although they are thought to function in part by acting as docking sites for transcription factors, which then activate appropriate target genes over long distances (Furlong and Levine, 2018).

Enhancers have been proposed to drive transcription at both the stages of initiation (Larke et al., 2021; Narita et al., 2021) and promoter-proximal pause release (Chen et al., 2018a; Core and Adelman, 2019). Live imaging has suggested that strong enhancers control the transcription cycle by increasing burst frequency rather than burst size (Fukaya et al., 2016). Roles for H3K27ac and factors such as BRD4 and Mediator have also been proposed, although the exact function of these complexes at enhancers has not been established (Donner et al., 2010; Galbraith et al., 2013; Hertweck et al., 2016; Kanno et al., 2014; Loven et al., 2013; Takahashi et al., 2011). Recent work in our lab has identified a subset of H3K79me2/3-marked Enhancer Elements (KEEs), which are dependent on this methylation for the maintenance of gene expression and enhancer-promoter interactions (Godfrey et al., 2019). What is not clear is the nature of the crosstalk between enhancers and promoters, or how the flow of information is controlled – in particular if the same or different complexes are important to drive the activity of both enhancers and promoters.

MLL-AF4, along with other MLL-FPs, is thought to drive leukemia by binding at the promoters of key oncogenes and upregulating their expression by the recruitment of a complex of elongation-associated factors, including the RNA Polymerase (RNAP) II-Associated Factor complex (PAF1C), Eleven Nineteen Leukemia (ENL/MLLT1), AF9, DOT1L, AFF4 and P-TEFb (Basu et al., 2020; Slany, 2020; Takahashi and Yokoyama, 2020). In many cases, MLL-FP binding spreads from the promoter into the gene body, associated with elevated levels of elongation factors and upregulated transcription (Kerry et al., 2017). Many target genes are also regulated by strong enhancers, and we and others have identified MLL-AF4 binding at intragenic and intergenic enhancers (Godfrey et al., 2021; Godfrey et al., 2017; Prange et al., 2017), but the significance of this behavior, and whether it is dependent on the same factors associated with MLL-AF4 at gene promoters, has not been established.

We sought to understand the role of MLL-AF4 in enhancer function, as a model to investigate mechanisms of aberrant enhancer activation in cancer. In order to characterise MLL-AF4 association with enhancers in the precise cellular context responsible for human disease, we adapted the low cell number high-throughput ChIPmentation method (Gustafsson et al., 2019) for primary samples, optimizing it for analysis of more weakly-associated non-histone chromatin proteins, an approach we call Transcription factor-OPtimized ChIPmentation (TOPmentation). This allowed us to generate coupled RNA-seq, ATAC-seq and chromatin profiles in four MLL-AF4 ALL patient samples.

We show that MLL-AF4 binding is a key driver of novel enhancer formation in human ALL and, furthermore, that MLL-AF4 is required to maintain multiple enhancer features, including proximity with target promoters. Many of the proteins colocalizing at MLL-AF4-bound enhancers are the same elongation factors which associate with MLL-AF4 at target genes, including ENL and PAF1C. We also identify a novel interaction between the MLL-AF4 complex and the histone chaperone Facilitates Chromatin Transcription (FACT). Chemical degradation of either PAF1 or the FACT component SSRP1 results in a loss of activity at many MLL-AF4-bound and -unbound enhancers, indicating that these factors are generally required for enhancer function. Together, this reveals an unexpected mechanism of oncogenicity of the MLL-AF4 fusion protein and argues that MLL-AF4 may promote transcription by assembling a large transcription elongation complex at both enhancers and promoters.

## Results

### MLL-AF4 binds at putative enhancers in primary patient samples

In order to characterize the chromatin environment in MLL-AF4 ALL patients, where the number of cells available for analysis is limiting, we developed TOPmentation, an optimization of the high-throughput ChIPmentation protocol (Gustafsson et al., 2019) to allow the capture of less tightly associated non-histone chromatin proteins and co-activators. By incorporating a double-fixation step, combining treatment with formaldehyde and disuccinimidyl glutarate, which has a longer linker, alongside additional optimizations to reduce background signal (see Methods), we were able to describe the chromatin binding profile of difficult-to-ChIP chromatin associated proteins from 100,000 cells. This includes MLL-AF4 itself as well as MLL-AF4 complex members such as ENL and PAF1 (Figure S1A). Building on existing ChIP-seq and Assay for Transposase-Accessible Chromatin (ATAC-seq) data (Godfrey et al., 2021; Godfrey et al., 2019; Harman et al., 2021; Kerry et al., 2017), we generated new data to assemble paired MLL-AF4 binding and chromatin profiles for leukemia cells from five MLL-AF4 ALL patients, including three childhood (chALL; 1-18 years old) and two infant (iALL; <12 months old) patients, with gene expression and chromatin accessibility data for four of these (Table S1). We performed TOPmentation and ChIP-seq for H3K4me1, H3K4me3, H3K27ac, H3K27me3, H3K79me2 and the N-terminus of MLL-AF4. The TOPmentation method generated highly comparable data to that produced by standard ChIP-seq using ∼100-fold fewer cells allowing the chromatin environment to be delineated even in patient samples where few primary leukemia cells are available (Figure S1B, C).

We and others have previously observed the binding of MLL-AF4 at enhancers as well as promoters in cell lines and individual patient samples (Godfrey et al., 2021; Godfrey et al., 2017; Prange et al., 2017). We therefore used TOPmentation to ask whether MLL-AF4 enhancer binding could be observed genome-wide in patient samples, by annotating MLL-AF4 peaks by distance to the nearest transcriptional start site (TSS). In all five patient samples analyzed, we saw a similar bimodal pattern of MLL-AF4 binding; while many MLL-AF4 peaks were present within 1 kb of the TSS, a large (but variable between samples) a proportion of peaks were more than 10 kb from the nearest TSS (Figure 1A). We observed a similar distribution in a range of *MLL*r ALL and AML cell lines (Figure S1D), indicating that non-promoter binding, possibly at enhancers, is a widespread property of MLL-FPs, and might suggest an alternative mechanism for regulating gene expression.

**Figure 1.**
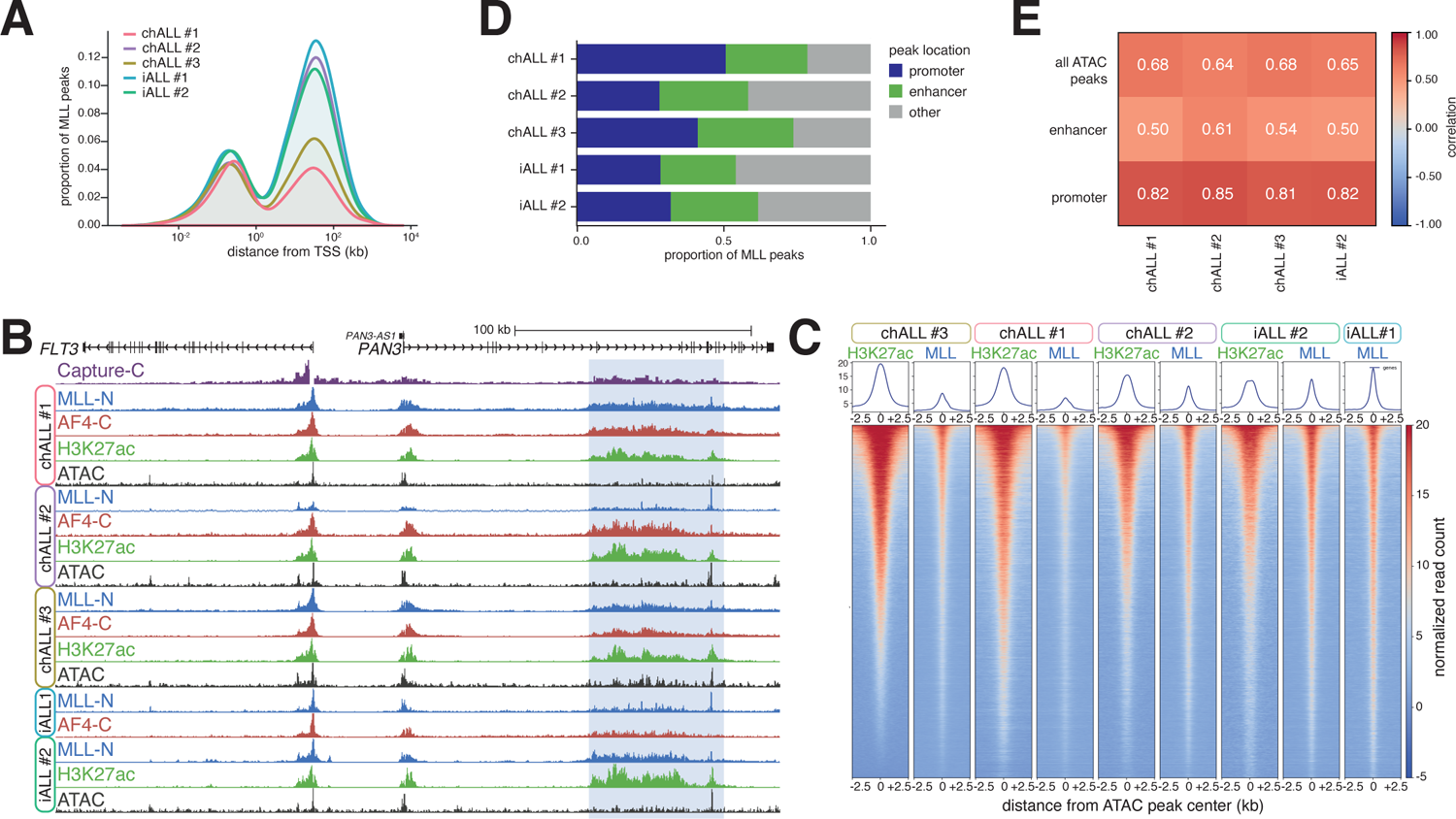
MLL-FP binding at enhancers is a common feature of cell lines and primary material. (A) Distribution of MLL peaks in five patient samples, relative to the nearest TSS. (B) ChIP-seq for MLL, AF4, H3K27ac, and ATAC-seq at *FLT3* and *PAN3* in the indicated patient samples. Capture-C from the TSS is shown for SEM cells. The MLL-AF4-bound enhancer within *PAN3* is highlighted in blue. (C) Heatmap of H3K27ac ChIP-seq and MLL ChIP-seq/TOPmentation signal in the indicated patient samples at enhancers. (D) Proportion of MLL peaks associated with promoters and enhancers in each patient sample. (E) Correlation of H3K27ac and MLL ChIP-seq/TOPmentation signal at promoters, enhancers and all ATAC peaks in the indicated patient samples.

Consistent with this, we found MLL-AF4-bound enhancers in proximity to a number of key oncogenes in patient samples, including *FLT3*, *MYC*, *CDK6* (Figure 1B, S1E) and *PROM1* (Godfrey et al., 2021), suggesting a potentially important role for these elements in driving and/or maintaining the leukemia. Notably, while many of these enhancers were associated with spreading of MLL-AF4 into the gene body from the promoter (Godfrey et al., 2021; Kerry et al., 2017), we also found MLL-AF4 binding at intergenic enhancers (e.g. *MYC*; Figure S1E, F), demonstrating that enhancer binding is not an indirect consequence of MLL-AF4 spreading from the promoter. Notably, we observed MLL-AF4 binding at an enhancer regulating the *ARID1B* gene in RS4;11 cells, where the enhancer is active, but not in SEM cells, where the enhancer is silent and does not interact with the *ARID1B* promoter (Figure S1G). Thus, in these specific examples and more generally there is a correlation between MLL-AF4 binding and enhancer activity.

The impact of MLL-AF4 binding on enhancer function has not been explored in detail. To address this, we used our patient data to systematically interrogate this behavior genome-wide. We identified putative enhancers by intersecting non-promoter ATAC-seq and H3K27ac peaks, and used MLL ChIP-seq/TOPmentation data to compare the binding of MLL-AF4 at these loci. MLL-AF4 was present at many of these enhancer regions (Figure 1C), and a substantial proportion of MLL-AF4 binding sites overlapped with enhancers in each patient (Figure 1D), suggesting that this is a common property of the fusion protein. We saw a strong correlation between levels of H3K27ac and MLL-AF4 binding at both promoters and enhancers (Figure 1E), indicating that MLL-AF4 associates with enhancer as well as promoter activity in primary ALL leukemia cells. This distribution may therefore play an important role in determining the biology of these leukemias.

### MLL-AF4 binding at enhancers is functionally relevant

Having established a genome-wide correlation between MLL-AF4 binding and enhancer activity, we asked whether the FP was required for enhancer function. We made use of the SEM cell line model, in which we have previously explored MLL-AF4 promoter function, where knockdown (KD) results in downregulation of a large number of genes (Kerry et al., 2017). To address the role of MLL-AF4 in enhancer activity, we intersected SEM enhancers and MLL-AF4 peaks, distinguishing those enhancers enriched (bound) or depleted (not bound) for MLL-AF4 (Figure 2A). Genes associated with an MLL-AF4-bound enhancer showed a stronger downregulation following MLL-AF4 KD than genes associated with an enhancer not bound by MLL-AF4 (Figure 2B), even when taking into account the presence/absence of MLL-AF4 at the promoter (Figure S2A). Further, a greater proportion of MLL-AF4 enhancer-associated genes were downregulated following MLL-AF4 KD (Figure 2B, S2B), consistent with the model that MLL-AF4 binding at these enhancers upregulates gene expression.

**Figure 2.**
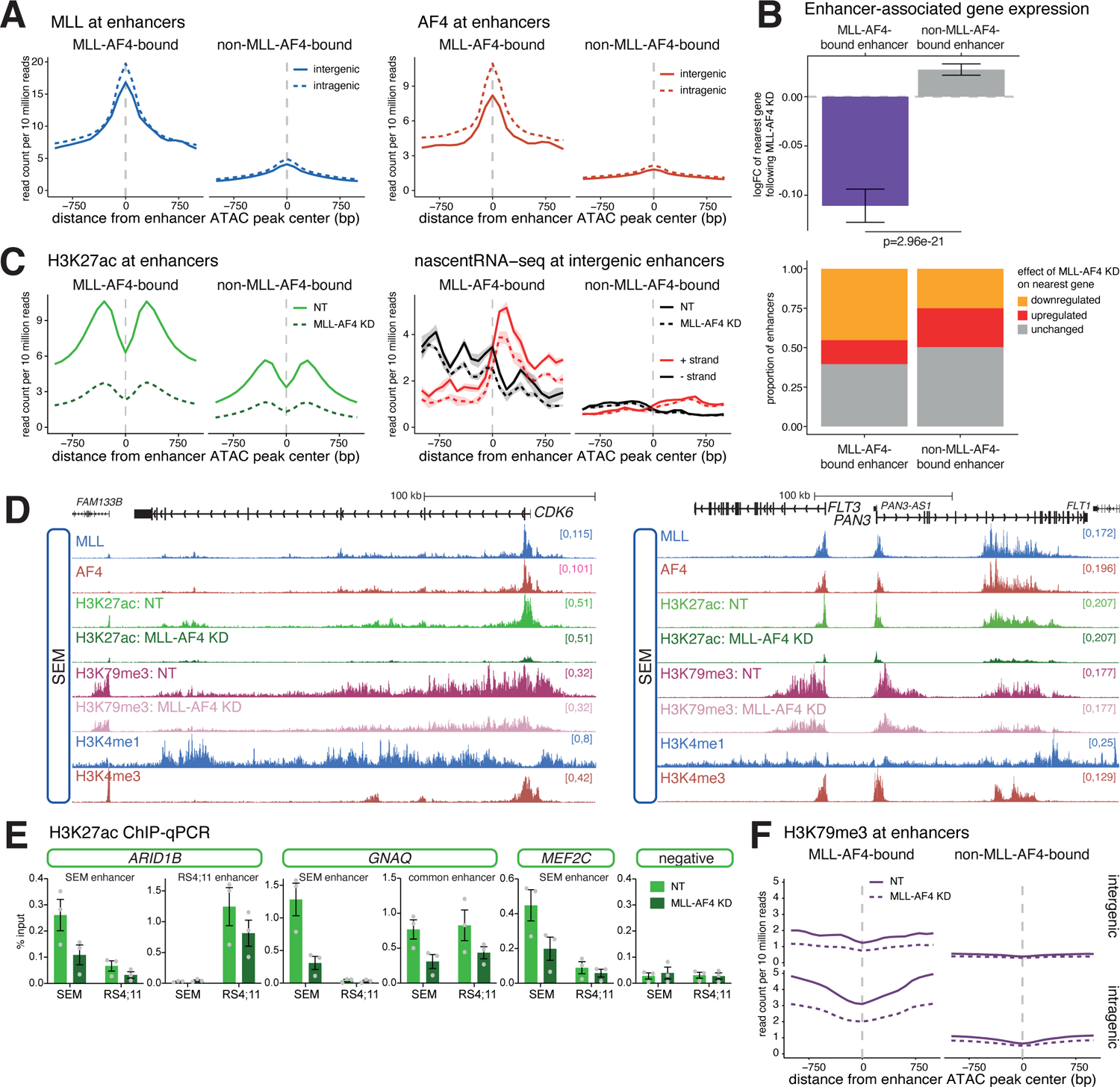
MLL-AF4 binding is required for the maintenance of enhancer signatures. (A) Mean distribution of MLL (*left*) and AF4 (*right*) at MLL-AF4-bound and non-MLL-AF4-bound intergenic (solid line) or intragenic (dashed line) enhancers in SEM cells. Plots are centered on ATAC-seq peaks found within enhancers. (B) *Upper*: Mean log-fold change in gene expression in SEM cells following MLL-AF4 KD, for genes associated with an MLL-AF4-bound enhancer or genes associated with an enhancer not bound by MLL-AF4. Statistical significance calculated using a Mann-Whitney U test. *Lower*: Proportion of enhancers associated with genes displaying each transcriptional response to MLL-AF4 KD. (C) *Left*: Mean distribution of H3K27ac at MLL-AF4-bound and non-MLL-AF4-bound enhancers, in SEM cells under control (NT) and MLL-AF4 KD conditions. *Right*: Mean distribution of strand-specific nascent RNA-seq (enhancer RNA) levels at MLL-AF4-bound and non-MLL-AF4-bound intergenic enhancers, under control (NT) and MLL-AF4 KD conditions. Lines represent mean, shading represents ± SEM, n=3. (D) Reference-normalized ChIP-seq for H3K27ac and H3K79me3 at *CDK6* and *FLT3* in SEM cells under control (NT) and MLL-AF4 KD conditions. ChIP-seq for MLL, AF4, H3K4me1 and H3K4me3 is shown for context. (E) ChIP-qPCR for H3K27ac at the indicated enhancer regions in SEM and RS4;11 cells, under control (NT) and MLL-AF4 KD conditions. Data are represented as mean ± SEM, n=3. (F) Mean distribution of H3K79me3 at MLL-AF4-bound and non-MLL-AF4-bound inter- and intragenic enhancers in SEM cells under control (NT) and MLL-AF4 KD conditions.

MLL-AF4 enrichment at enhancers correlated with elevated H3K27ac and enhancer RNA (eRNA) transcription (measured by nascent RNA-seq), suggesting that MLL-AF4 is associated with high levels of enhancer activity (Figure 2C). KD of MLL-AF4 resulted in a general reduction in enhancer acetylation, but there was a much larger decrease at MLL-AF4-bound enhancers (Figure 2C, D), suggesting this is a direct consequence of loss of MLL-AF4 binding. The effect was confirmed by ChIP-qPCR in both SEM and RS4;11 cells at common and cell line-specific enhancers (Figure 2E, S2D). We also observed a clear reduction in eRNA transcription from MLL-AF4-bound enhancers following MLL-AF4 KD (Figure 2C, right), arguing for a loss of activity. Together, these results strongly indicate a functional role for MLL-AF4 in enhancer activity.

We have previously shown that H3K79me2/3 is required to maintain the activity of a subset of enhancers, termed H3K79me2/3-marked enhancer elements (KEEs) (Godfrey et al., 2019). MLL-AF4 is associated with elevated H3K79me2/3 (Krivtsov et al., 2008; Rice et al., 2021), and we observed an enrichment for H3K79me3 at both intragenic and intergenic enhancers bound by MLL-AF4 (Figure 2F). Indeed, many KEEs are bound by MLL-AF4 (Figure S2E). This mark was depleted following MLL-AF4 KD (Figure 2F), suggesting that MLL-AF4 may be responsible for generating a subset of KEEs in SEM cells. For example, the MLL-AF4-bound enhancer elements within *CDK6*, *FLT3* and *ARID1B* showed a strong depletion of H3K27ac and H3K79me3 following MLL-AF4 KD (Figure 2D, S2C), associated with downregulation of gene transcription (Figure S2F).

### MLL-AF4 recruits elongation factors to active enhancers

To understand how MLL-AF4 promotes enhancer activity, we asked whether the fusion protein recruits core components of the promoter-associated MLL-AF4 complex to enhancers. MLL-AF4 is known to interact with a core set of factors that includes DOT1L, MENIN, ENL and a large number of other proteins (Biswas et al., 2011; Lin et al., 2010; Yokoyama et al., 2010). The elevated levels of H3K79me3 (Figure 2F), commonly associated with MLL-AF4 binding, suggest the assembly of a functional complex of factors at enhancers, including DOT1L. In addition, MLL-AF4 enhancers were enriched for binding of MENIN and ENL (Figure 3A). Levels of these proteins were elevated at both intergenic and intragenic enhancers, indicating that the presence of these factors was not an indirect consequence of increased gene transcription or spreading from promoters (Figure 3A).

**Figure 3.**
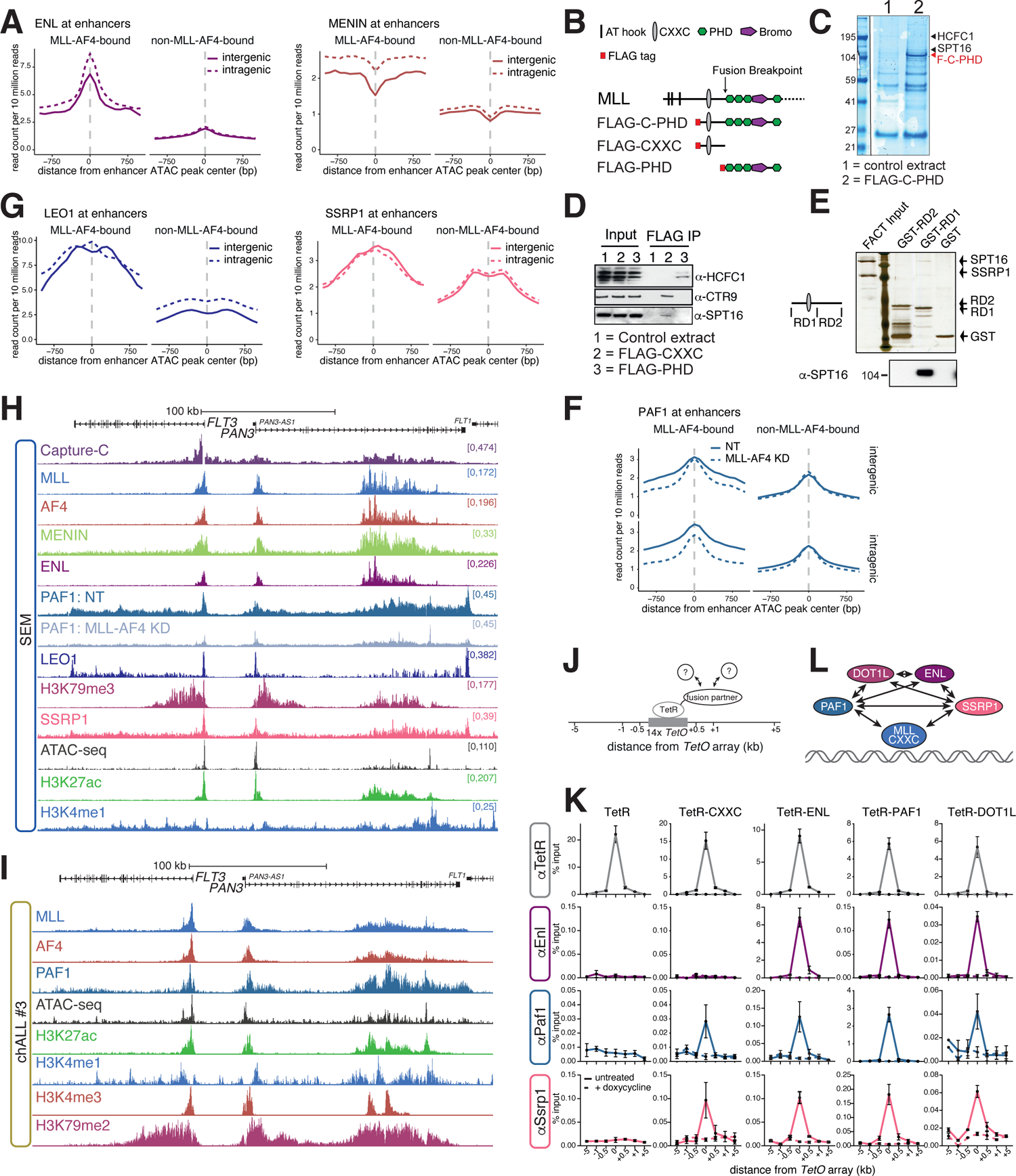
MLL-AF4 binding recruits transcription elongation factors to enhancers. (A) Mean distribution of ENL and MENIN at MLL-AF4-bound and non-MLL-AF4-bound intergenic (solid line) and intragenic (dashed line) enhancers in SEM cells. (B) Schematic of wild-type N-terminal MLL structure, showing the domains used for immunoprecipitation. (C) Colloidal Blue-stained gel of control HEK 293 nuclear extracts (1) or HEK 293 nuclear extracts expressing the FLAG-C-PHD construct (2), immunoprecipitated with anti-FLAG antibody. Gel lanes were sliced and subjected to mass spectrometry (see Methods). Regions where the proteins HCFC1 and FACT complex component SPT16 were identified are indicated by arrowheads. The red arrowhead indicates the position of the FLAG-C-PHD protein. (D) Immunoblots for HCFC1, CTR9 (PAF1C component) and SPT16 following anti-FLAG immunoprecipitation of HEK 293 cell lysates expressing the indicated FLAG-tagged MLL domains. Representative of three experiments. (E) Silver-stained gel after affinity purification of GST-tagged MLL RD1 and RD2 domains following incubation with purified SPT16 and SSRP1 (FACT). Lower panel shows immunoblot for SPT16. Representative of two experiments. (F) Mean distribution of PAF1 at MLL-AF4-bound and non-MLL-AF4-bound intragenic and intergenic enhancers in SEM cells under control (NT; solid line) and MLL-AF4 KD (dashed line) conditions. (G) Mean distribution of PAF1C component LEO1 and FACT component SSRP1 at MLL-AF4-bound and non-MLL-AF4-bound intergenic (solid line) and intragenic (dashed line) enhancers in SEM cells. (H) ChIP-seq, ATAC-seq and Capture-C at the *FLT3* locus in SEM cells. Reference-normalized ChIP-seq for PAF1 in SEM cells under control (NT) and MLL-AF4 KD conditions. The Capture-C viewpoint is the *FLT3* TSS. (I) TOPmentation and ATAC-seq at the *FLT3* locus in chALL patient #3. (J) Schematic showing the principle behind the TetR recruitment system. (K) ChIP-qPCR for the indicated proteins (left) at the *TetO* array inserted into mESCs expressing the indicated TetR fusion proteins (top). Dashed line shows ChIP-qPCR in cells treated with doxycycline for 6h. Data are represented as mean ± SEM, n≥3. (L) Model indicating direct or indirect in vivo interactions demonstrated in (K).

In addition to the MLL-AF4 core complex, we looked at additional MLL-AF4-interacting proteins that may be recruited to enhancers. For example, PAF1C is a known co-activator (Kim et al., 2010) that interacts directly with the CXXC domain of MLL (Milne et al., 2010; Muntean et al., 2010). By screening a previously unpublished MLL mass spectrometry dataset for additional co-activators, we also identified the histone chaperone FACT as a potential MLL-interacting protein (Figure 3B, C, Table S2). Both PAF1C and FACT are known to travel with RNAPII and enhance transcription (Francette et al., 2021; Kim et al., 2010; Orphanides et al., 1998; Wang et al., 2021), so we wanted to determine if MLL-AF4 binding was responsible for increased enrichment of these factors. To validate the putative interaction between MLL-AF4 and FACT, we performed immunoprecipitation (Figure 3B, D) and GST pulldown experiments (Figure 3E) and showed that purified FACT complex interacts directly with the CXXC domain-containing region of MLL. Taken together, these data suggest that FACT is a novel component of both wild type and MLL-FP complexes, binding directly to the CXXC domain of MLL.

We observed elevated levels of components of PAF1C (PAF1 and LEO1) and FACT (SSRP1) at both intergenic and intragenic MLL-AF4-bound enhancers, suggesting that MLL-AF4 may recruit PAF1C and FACT to enhancers (Figure 3F, G, H, S3A). To confirm that this also occurred in primary ALL cells we used PAF1 and H3K79me2 TOPmentation (Fig 3I, S3B). MLL-AF4 binding correlated strongly with PAF1 levels at both promoters and enhancers of the patient sample (Figure S3C), for example at *ARID1B* and *FLT3*/*PAN3* (Figure 3I, S3B). MLL-AF4 appears to be required for PAF1 enrichment, as MLL-AF4 KD reduced levels of PAF1 at both intergenic and intragenic MLL-AF4-bound enhancers, but not at non-MLL-AF4-bound enhancers, arguing for a direct stabilization of the complex at these loci (Figure 3F). These results suggest that the fusion protein may promote enhancer activity by assembling the same transcription-promoting complex as at promoters, both in cell line model systems and in primary human leukemia cells.

To understand how FACT and other proteins may associate with MLL-AF4 enhancers, we turned to a targeted recruitment system to test in vivo interactions (Figure 3J). We generated mESC lines expressing individual components of the MLL-AF4 complex fused to TetR, which allows them to bind stably at an array of *TetO* repeats inserted into the mouse genome (Blackledge et al., 2014). We used ChIP-qPCR to assess the ability of these proteins to recruit other factors to the *TetO* array, indicating an in vivo interaction (Figure 3K). Consistent with biochemical analyses (Biswas et al., 2011; He et al., 2011; He et al., 2010; Leach et al., 2013; Lin et al., 2010; Milne et al., 2010; Mohan et al., 2010; Mueller et al., 2007; Muntean et al., 2010; Sobhian et al., 2010; Yokoyama et al., 2010) we observed multiple reciprocal interactions between ENL, PAF1 and DOT1L (Figure 3K). As previously reported (Milne et al., 2010; Muntean et al., 2010), the CXXC domain of MLL was able to weakly recruit Paf1, indicating that PAF1C may be localized to MLL-AF4-bound loci by interaction with both the fusion protein and other complex components. Strikingly, we saw a similar effect with FACT, where the MLL CXXC domain, ENL, PAF1 and DOT1L were all able to recruit Ssrp1 to the *TetO* array (Figure 3K, bottom row). Consistent with this, FACT has previously been demonstrated to interact with Paf1 in yeast (Krogan et al., 2002; Squazzo et al., 2002). We also verified that the FACT:MLL-CXXC domain interaction occurs specifically through the CXXC-RD1 domain (Figure S3D). Together, this argues that colocalization of FACT and the elongation machinery with MLL-AF4 may be achieved via multivalent interactions with multiple components of the complex (Figure 3L).

### PAF1C and FACT are required for the activity of enhancers, independent of MLL-AF4 binding

In order to assess the role of PAF1C and FACT in enhancer function, we generated SEM degron cell lines where endogenous PAF1 or SSRP1 was tagged with the FKBP12^F36V^ domain (Nabet et al., 2018). Treatment of cells with the small molecule dTAG-13 results in rapid and dramatic reduction in protein levels (Figure 4A, S4A). We assessed the effect of protein degradation on gene expression by reference-normalized transient transcriptome sequencing (TT-seq) (Schwalb et al., 2016). This revealed a significant downregulation of transcription at the vast majority of expressed genes (Figure S4B). From these experiments it is not possible to determine whether loss of each factor primarily affected transcription elongation or initiation. However, degradation of PAF1 had a stronger effect on detectable transcripts towards the 3’ end of the gene, whereas SSRP1 loss appeared to reduce transcripts in a more uniform pattern, potentially indicating a stronger effect on transcription initiation (Figure S4C).

**Figure 4.**
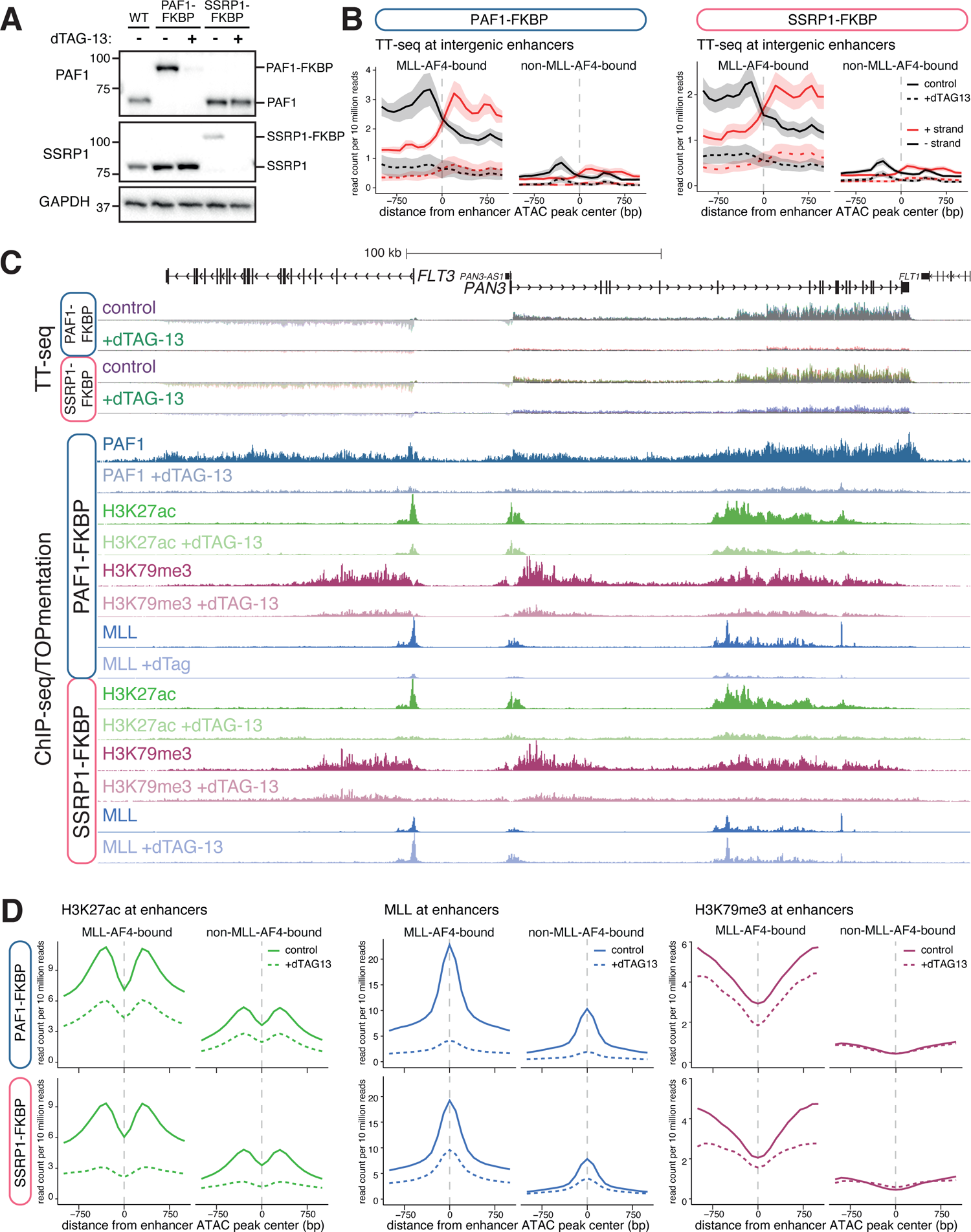
PAF1 and SSRP1 are required for activity of MLL-AF4-bound and non-MLL-AF4-bound enhancers. (A) Western blot for PAF1 or SSRP1 in wild-type (WT), PAF1-FKBP or SSRP1-FKBP cells, with (+) or without (-) addition of 0.5 µM dTAG-13 for 24h. Blots are representative of three replicates. (B) Mean distribution of strand-specific TT-seq (eRNA) levels at MLL-AF4-bound and non-MLL-AF4-bound intergenic enhancers, in PAF1-FKBP or SSRP1-FKBP cell lines under control (untreated) and dTAG13-treated conditions. Lines represent mean, shading represents ± SEM, n=3. (C) TT-seq, reference-normalized ChIP-seq and TOPmentation at the *FLT3* locus in PAF1-FKBP and SSRP1-FKBP SEM cells, with or without the addition of dTAG-13. (D) Mean distribution of H3K27ac, MLL and H3K79me3 at MLL-AF4-bound and non-MLL-AF4-bound enhancers, in PAF1-FKBP (above) or SSRP1-FKBP (below) cell lines under control (untreated) and dTAG13-treated conditions.

Genes associated with either an MLL-AF4-bound or non-MLL-AF4-bound enhancer were strongly downregulated by PAF1 or SSRP1 degradation (Figure S4D). To understand the role of PAF1C and FACT at enhancers, we analyzed eRNA transcription. As with MLL-AF4 KD (Figure 2C), degradation of either PAF1 or SSRP1 resulted in a dramatic reduction in eRNA levels at MLL-AF4-bound enhancers (Figure 4B), suggesting a key role for these factors in this enhancer activity. This effect was particularly striking at the intragenic *FLT3*/*PAN3* enhancer (Figure 4C). However, in contrast to MLL-AF4 KD (Figure 2C), we also observed a reduction in eRNA transcription at non-MLL-AF4-bound enhancers (Figure 4B), indicating that PAF1C and FACT have a more general role at enhancers, beyond their association with MLL-AF4. For example, at *LMO4*, we observed a reduction in enhancer activity in the absence of MLL-AF4 (Figure S4E). To confirm this effect, we analyzed H3K27ac levels at enhancers following degradation of PAF1 or SSRP1. As with eRNA transcription, levels of acetylation were reduced at both MLL-AF4-bound and not bound enhancers (Figure 4C, D, S4E). These decreases were observed at intergenic as well as intragenic enhancers, arguing that this is not an indirect effect of loss of gene transcription (Figure S4F).

Given the multivalent interactions we observed between PAF1C, FACT and members of the MLL-AF4 complex, we asked whether PAF1C and FACT might themselves stabilize binding of MLL-AF4 at enhancers. Indeed, we observed a decrease in MLL binding at enhancers following loss of PAF1 or SSRP1, with a particularly striking reduction associated with PAF1 degradation (Figure 4D). H3K79me3 levels were also reduced at MLL-AF4 enhancers, suggesting a loss of DOT1L association/activity (Figure 4D, S4F). Thus, we hypothesize that PAF1C and FACT may contribute to enhancer function both directly (by promoting eRNA transcription) and indirectly (via MLL-AF4 complex stabilization).

### Enhancer-promoter contacts are differentially dependent on MLL-AF4, PAF1C and FACT

A key feature of most enhancers is a high frequency of interaction with target gene promoters. We hypothesized that MLL-AF4 and the complex of proteins it assembles at enhancers may play a role in driving these interactions. We used Next Generation Capture-C (Davies et al., 2016; Downes et al., 2022) to test whether physical contact between MLL-AF4 enhancers and promoters was dependent on the presence of the fusion protein. Using promoter-specific capture probes, we observed a striking correlation between MLL-AF4 binding at enhancers and promoter interaction frequency (Figure 5A, S5A, purple shading). This is particularly clear when comparing the interaction profiles at the same gene in SEM and RS4;11 cells; in each cell type, enhancer interaction mirrors the binding profile of MLL-AF4 (Figure S5B).

**Figure 5.**
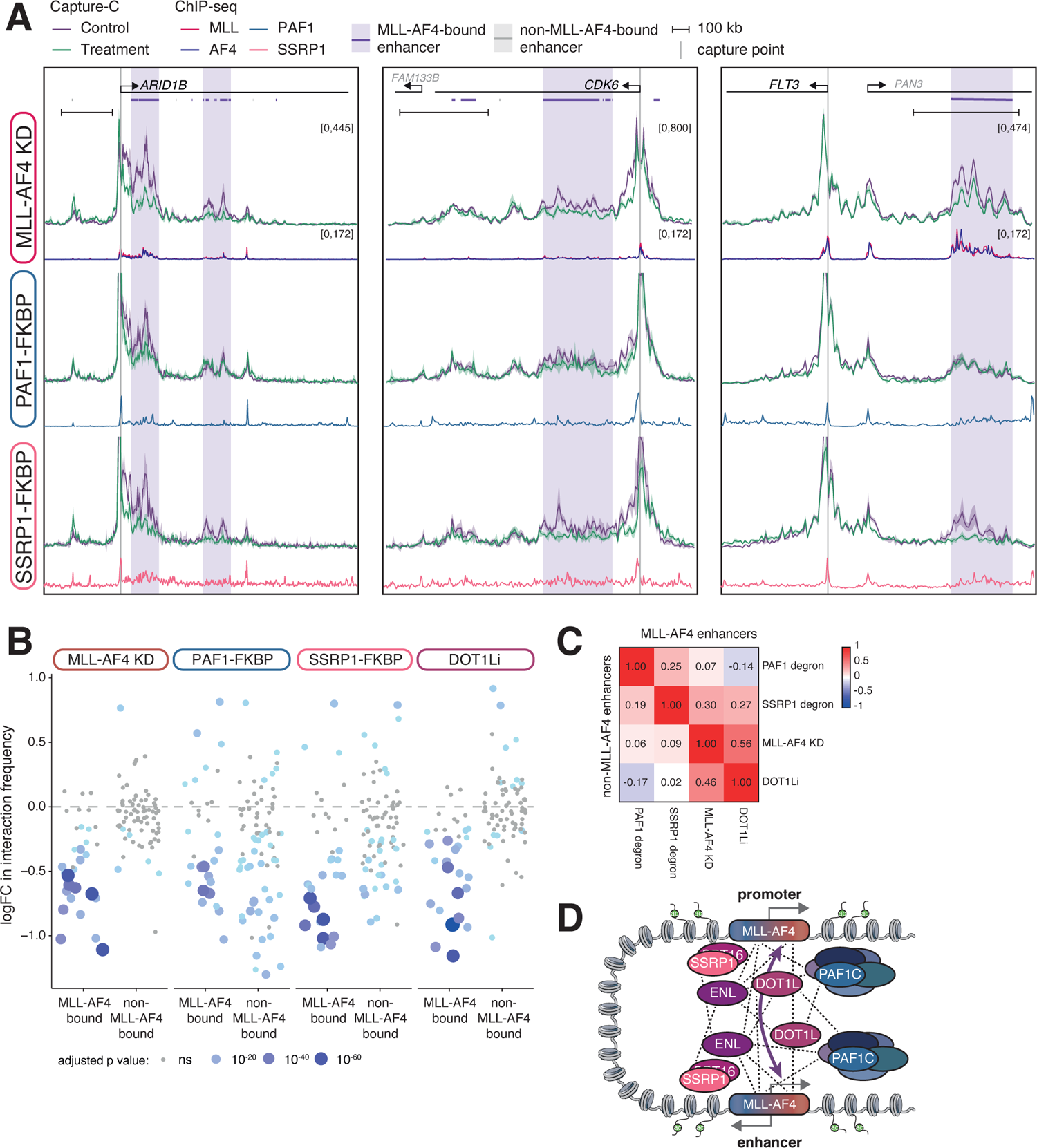
MLL-AF4 binding is necessary to maintain enhancer-promoter interactions. (A) Capture-C from the promoters of *ARID1B*, *CDK6*, and *FLT3* in SEM cells under control (purple) and MLL-AF4 KD (green) conditions (*upper*) or in PAF1-FKBP or SSRP1-FKBP cell lines under control (purple) and dTAG13-treated (green) conditions. Lines represent mean, shading represents ± SEM, n=3. ChIP-seq traces for MLL, AF4, PAF1 and SSRP1 are shown, along with bioinformatically-annotated MLL-AF4-bound and -unbound enhancers. (B) Statistical analysis of the changes in Capture-C interaction frequency between promoters and MLL-AF4-bound or -unbound enhancers, following the indicated treatments. Size and color of the dot is proportional to the significance of the change. n=3; ns: adjusted p value ≥0.05; Mann–Whitney U test. DOT1Li Capture-C data taken from Godfrey et al 2019. (C) Correlation of changes in Capture-C interaction frequency between promoters and MLL-AF4-bound (upper triangle) or -unbound (lower triangle) enhancers, comparing the indicated treatments. (D) Model for the role of MLL-AF4 at enhancers in driving interaction with and transcription of target genes, recruiting a complex of transcription-associated proteins. Dashed lines indicate the network of protein-protein interactions.

Strikingly, MLL-AF4 KD in SEM cells significantly reduced the frequency of interaction of promoters with MLL-AF4-bound enhancers (Figure 5A, S5A), but not enhancers lacking MLL-AF4 (Figure S5C). We analyzed a panel of 32 genes, including genes associated with MLL-AF4-bound or non-MLL-AF4-bound enhancers (Table S3), and found a clear and specific effect of MLL-AF4 KD on promoter interactions with MLL-AF4-bound enhancers (Figure 5B, S5D). This argues that MLL-AF4 binding at the enhancer is required to drive and/or stabilize contact with the promoter. Together with the effects of MLL-AF4 KD on other enhancer features, this indicates a key role for MLL-AF4 in the function of directly bound enhancers.

We next asked whether the recruitment of elongation factors to enhancers by MLL-AF4 could provide a mechanism by which the FP achieves enhancer-promoter contact. We used our PAF1/SSRP1 degron cell lines to test whether these enhancer-promoter interactions were dependent on the presence of PAF1C or FACT. While the effect was less pronounced than with MLL-AF4 KD, many enhancer-promoter interactions were sensitive to degradation of PAF1 or SSRP1 (Figure 5A, B, S5A). However, in contrast to MLL-AF4 KD, there was less bias towards MLL-AF4-bound enhancers, with reductions in promoter interactions also observed with many non-MLL-AF4-bound enhancers (Figure 5B, S5C). This is consistent with the observation that eRNA and H3K27ac levels at both MLL-AF4 and non-MLL-AF4-bound enhancers were sensitive to loss of PAF1 and SSRP1 (Figure 4B, D).

MLL-AF4-bound enhancers are marked with H3K79me3 (Figure 2F), and the majority of these enhancers are annotated as KEEs (Figure S2E). Consistent with this, promoter interaction frequency was much more sensitive to MLL-AF4 KD at KEEs compared to non-KEEs (Figure S5D). We have previously shown that KEEs are sensitive to chemical disruption of H3K79me2/3 by DOT1L inhibition, resulting in a loss of enhancer-promoter contact (Godfrey et al., 2019). KD of MLL-AF4 resulted in reduced H3K79me3 levels at enhancers (Figure 4C), suggesting that DOT1L enrichment may provide a potential mechanism for stabilizing enhancer-promoter interactions. A direct comparison of our MLL-AF4 KD Capture-C with previously published Capture-C at the same genes following DOT1L inhibition (Godfrey et al., 2019), revealed a positive correlation (R = 0.56) in the change in interaction frequency between promoters and MLL-AF4 enhancers (Figure 5C, S5E), indicating that loss of H3K79me2/3 partly reproduces the effect of MLL-AF4 depletion at these loci. We observed a much weaker correlation between MLL-AF4 KD and PAF1 or SSRP1 degradation (Figure 5C, S5E), suggesting that these factors may have specific roles at a subset of MLL-AF4-bound and non-MLL-AF4-bound enhancers. From this, we conclude that MLL-AF4 may drive enhancer activity by recruiting transcriptional machinery to upregulate levels of H3K79 methylation. Transcription elongation factors such as PAF1C and FACT are also recruited by MLL-AF4, with a role in maintaining enhancer-promoter interactions (Figure 5D), but they display a more general role at enhancers and their activity is not specifically limited to MLL-AF4 enhancers.

## Discussion

Enhancer activation is a common mechanism for gene upregulation in cancer (Hnisz et al., 2015; Loven et al., 2013; Sur and Taipale, 2016). We and others have identified key enhancers in *MLL*r leukemias (Godfrey et al., 2021; Godfrey et al., 2019; Godfrey et al., 2017; Prange et al., 2017; Tzelepis et al., 2018), but it has been unclear what drives aberrant enhancer activity in these cells. We show here a direct requirement for MLL-AF4 at enhancers, which functions by assembling a complex of factors to drive enhancer activity, including ENL, PAF1C, DOT1L and FACT. MLL-AF4 activates a network of TFs, directing a program of leukemic transcription (Harman et al., 2021) and we note that it is likely that some enhancers are also activated indirectly via the activation of TFs within this network.

MLL-AF4-bound enhancers are associated with high levels of H3K27ac and eRNA transcription, and MLL-AF4 KD results in a reduction of both features. Strikingly, there is also a significant reduction in interaction frequency between the enhancer and promoter, arguing that MLL-AF4 is required to establish or maintain this proximity. This is unlikely to be an indirect consequence of the loss of transcription, as we have previously shown that inhibition or degradation of BRD4 significantly downregulates gene expression without affecting enhancer-promoter contacts (Crump et al., 2021). It is unclear how these effects are mediated, but the presence of the same complex of MLL-AF4 associated proteins at enhancers and gene promoters, including ENL, PAF1C, DOT1L and FACT, suggests that they may have a role in stimulating transcription from these distal loci. Indeed, chemical degradation of PAF1 or FACT component SSRP1 dramatically reduced not only gene transcription (as expected for elongation factors), but also eRNA transcription and other enhancer features, suggesting additional, enhancer specific, roles for these complexes.

Remarkably, loss of PAF1C or FACT also resulted in a reduction in the frequency of interaction between specific enhancer and promoter loci, indicating roles for these factors in 3D contacts. Degradation of PAF1 or SSRP1 produced distinct, but overlapping, effects on enhancer activity compared to MLL-AF4 KD. This suggests that they have roles at enhancers independent of MLL-AF4, and beyond *MLL*r leukemia. MLL-AF4 may promote the activity of some enhancers by recruiting these factors to them. It is possible that context plays an important role in determining which enhancers are dependent on PAF1C and FACT. In contrast, many of the effects of MLL-AF4 KD on enhancer activity were replicated by DOT1L inhibition (Godfrey et al., 2019), suggesting that this activity is key to function at many MLL-AF4-bound enhancers. Thus, it is likely that MLL-AF4 acts via the localization of multiple factors to enhancers to drive transcriptional upregulation.

The precise function of PAF1C in transcription is unclear. In vitro transcription models demonstrated a role in transcription elongation (Kim et al., 2010). However, while most in vivo experiments suggest it may promote elongation by regulating promoter-proximal pause release (Hou et al., 2019; Lu et al., 2016; Wu et al., 2014; Yu et al., 2015), others have argued for suppressive activity (Bai et al., 2010; Chen et al., 2015; Chen et al., 2017). Recent work has identified distinct requirements for different PAF1C components in stimulating RNAP II activity (Zumer et al., 2021). In contrast to our work, the authors found that degradation of the PAF1 subunit only weakly affected RNA synthesis, which may reflect the shorter treatment time used (Zumer et al., 2021). PAF1C binding has previously been observed at enhancers (Chen et al., 2017; Ding et al., 2021), but it is not known how it localizes at these sites. We show here that MLL-AF4 localizes PAF1C to enhancers via multiple direct and indirect interactions in a co-dependent manner, consistent with a role in promoting transcription of target genes. It is unclear how PAF1C is recruited to enhancers independent of MLL-AF4, although there may be a role for wild-type MLL. Our data complement a recent study showing a strong correlation between PAF1C binding and enhancer activity (Ding et al., 2021), arguing for a role in promoting transcription.

The FACT complex functions in transcription by displacing histone H2A/H2B dimers from nucleosomes, promoting access for RNAP II transcription (Chen et al., 2018b; Orphanides et al., 1998). Consistent with the effects we observe following SSRP1 degradation, depletion of FACT in yeast reduces both transcription and RNAP II assembly at promoters (Petrenko et al., 2019). However, the role of FACT at enhancers is unclear, with some studies arguing it may suppress expression of target promoters (Ferri et al., 2019), although our data argue for a role in promoting gene activation. As has been proposed for PAF1C (Ding et al., 2021), FACT may perform a similar function at enhancers as in the gene body, facilitating RNAP II transcription, in this case for the production of eRNAs. An alternative but not exclusive possibility is that FACT disruption of nucleosome structure at enhancers aids in the binding of TFs, which in turn drive enhancer features, for example stabilization of enhancer-promoter interactions.

To understand how MLL-AF4 assembles a transcriptional complex at promoters and enhancers, we used the TetR binding assay to identify in vivo interactions. DOT1L and H3K79me2/3 are known to be associated with MLL-AF4 target genes, but how they are recruited is unclear. ENL/AF9 can interact with DOT1L via their ANC homology domain (AHD) (Leach et al., 2013; Li et al., 2014; Mueller et al., 2007). However, this domain is also the site of interaction with the AF4 fusion partner, placing MLL-AF4 and DOT1L in mutually exclusive complexes (Biswas et al., 2011; Leach et al., 2013; Yokoyama et al., 2010). Thus, ENL/AF9 alone cannot mediate the MLL-AF4/DOT1L interaction. Our data suggest an alternative mechanism, demonstrating that DOT1L is also able to interact with PAF1C, consistent with the fact that Paf1 is required for H3K79 methylation in yeast (Krogan et al., 2003; Wood et al., 2003). We also confirmed previous findings that PAF1C binds to MLL via the MLL CXXC domain (Milne et al., 2010; Muntean et al., 2010) and ENL via its YEATS domain (He et al., 2011; Hetzner et al., 2018), indicating two mechanisms by which PAF1C could associate with the MLL-AF4 complex. The ability of both the MLL and AF4 portions of the FP to interact with many of these proteins may explain the strong enrichment of factors observed at MLL-AF4 binding sites. Thus, a network of multivalent interactions may be responsible for assembling and stabilizing the complex of proteins associated with MLL-AF4 at chromatin. The presence of multiple interactions could allow for the concentration of factors at a locus without requiring continuous binding. We identified a new component of this complex, the histone chaperone FACT, which interacts with MLL, ENL, PAF1C and DOT1L in vivo via similar multivalent contacts. FACT has previously been found to interact with other TFs, for example Oct4 (Ding et al., 2012; Pardo et al., 2010) and TIF1γ/TRIM33 (Bai et al., 2010; Charles Richard et al., 2016; Ferri et al., 2019), suggesting that this may be a common mechanism to promote transcription. Interestingly, *MLL*r leukemias have been shown to be sensitive to FACT inhibition (Somers et al., 2020).

How might MLL-AF4 facilitate the physical proximity of enhancer and promoter? As noted, the fusion protein is associated with a large complex of proteins at both sites, involving numerous multivalent interactions. The idea that multiple weak interactions might drive complex assembly, stability, and ultimately enhancer function is consistent with recent models proposing the assembly of phase condensates as being drivers of enhancer activity (Boija et al., 2018; Cho et al., 2018; Guo et al., 2019; Hnisz et al., 2017; Sabari et al., 2018; Shrinivas et al., 2019; Zamudio et al., 2019). For MLL-AF4, multivalent interactions include the direct interaction between the AF4 component of the fusion protein and the C-terminus of wild-type AF4, or potentially other MLL-AF4 molecules (Biswas et al., 2011; Lin et al., 2010; Yokoyama et al., 2010). We have previously proposed that these interactions drive spreading of MLL-AF4 in cis from the promoter into the gene body (Kerry et al., 2017), and our Capture-C data here suggest that they may also be sufficient for trans interactions between MLL-AF4-bound at the enhancer and promoter (Figure 5D). Thus, MLL-AF4 and the complex of proteins with which it associates may act as a bridge between enhancer and promoter, possibly acting in concert with cohesin and CTCF, which are found at many of these loci (Crump et al., 2021). Degradation of PAF1 or SSRP1 disrupts MLL-AF4 binding, and DOT1L inhibition leads to loss of DOT1L binding to chromatin (Wu et al., 2021), suggesting that complete complex disassembly may explain the loss of enhancer-promoter interactions observed when individual components are either inhibited or degraded.

Finally, although model systems are essential to fully define the precise mechanisms that underpin the oncogenic behavior of MLL-AF4, it is important to understand how these systems reflect the biochemical events taking place in primary ALL cells in patients. Characterizing the distribution of MLL-AF4 binding and the chromatin profile of MLL-AF4-bound regions has been severely hampered by the limitations of experimental methodology applicable to the small numbers of cells available in patient samples. Although several key datasets have been published exploring the transcriptional profiles of MLL-AF4 ALL in primary patient samples (Agraz-Doblas et al., 2019; Andersson et al., 2015), there are few examples of chromatin analysis, often only in a single patient in the absence of coupled transcription data (Godfrey et al., 2021; Harman et al., 2021; Janssens et al., 2021; Kerry et al., 2017), limiting a detailed exploration of what drives gene expression in these leukemias, as opposed to cell line models. For example, CUT&Tag has recently been employed as an approach for generating genome-wide binding profiles in lower cell numbers (Janssens et al., 2021). In order to study the mechanism of MLL-AF4 activity in a disease-relevant context, we have developed an optimized ChIPmentation technique, TOPmentation. This allows analysis of difficult-to-ChIP factors in low cell number samples, enabling the simultaneous description of multiple histone modifications and transcription factors from a single sample, alongside paired ATAC-seq and RNA-seq. With this comprehensive profiling, we demonstrated MLL-AF4 binding at enhancers in patients, and strong colocalization with PAF1 at these enhancers. We believe that this dataset will be a valuable resource for research into the transcriptional regulation of *MLL*r leukemia.

## Methods

### Patient samples

Infant (<1 year old) and childhood ALL samples were obtained from Blood Cancer UK Childhood Leukaemia Cell Bank, UK (REC: 16/SW/0219). All ALL samples were anonymized at source, assigned a unique study number and linked. Cryopreserved samples were thawed, and immediately used for RNA extraction/ATAC analysis or fixed for ChIP-seq/TOPmentation.

### Cell culture and RNA interference

SEM cells (Greil et al., 1994) were purchased from DSMZ (www.cell-lines.de) and cultured in Iscove’s Modified Dulbecco’s Medium (IMDM) with 10% fetal calf serum (FCS, Gibco) and Glutamax (ThermoFisher Scientific). RS4;11 cells were purchased from ATCC (www.lgcstandards-atcc.org) and cultured in RPMI 1640 with 10% FCS and Glutamax. Mouse ES cells with a TetO-array (TOT2N mESC) were kindly provided by Dr. Rob Klose (University of Oxford) and were grown in DMEM supplemented by with 10% FCS, NEAA, Glutamax, LIF and β-mercaptoethanol. All cell lines were confirmed free from mycoplasma contamination.

siRNA knockdown of MLL-AF4 was conducted as previously described (Kerry et al., 2017), using the following sequences: **NT siRNA**: sense AAAAGCUGACCUUCUCCAAUG; antisense CAUUGGAGAAGGUCAGCUUUUCU. **SEM MLL-AF4 KD siRNA**: sense AAGAAAAGCAGACCUACUCCA; antisense UGGAGUAGGUCUGCUUUUCUUUU. **RS4;11 MLL-AF4 KD siRNA**: sense ACUUUAAGCAGACCUACUCCA; antisense UGGAGUAGGUCUGCUUAAAGUCC.

### Chromatin immunoprecipitation

Chromatin immunoprecipitation was conducted as previously described (Godfrey et al., 2019; Kerry et al., 2017). Briefly, 10^7^ cells were single-fixed (1% formaldehyde for 10 min) for histone modifications or double-fixed (2 mM disuccinimidyl glutarate for 30 min, then 1% formaldehyde for 30 min) for transcription/chromatin factors, then lysed with 120 µl SDS lysis buffer (10 mM Tris-HCl pH 8.0, 1 mM EDTA, 1% SDS) and sonicated using a Covaris ME220 (Woburn, MA) to generate 200-300 bp fragments. Insoluble material was pelleted, then the supernatant was diluted 10x and pre-cleared for 30 min at 4 °C with rotation using 5 µl protein A- and G-coupled dynabeads (ThermoFisher Scientific). An input sample was taken, then 2 µg antibody added to the sample before incubation overnight at 4 °C with rotation. Antibodies used are detailed in Table S4. Protein A- and G-coupled dynabeads were used to isolate antibody-chromatin complexes, after which the beads were washed three times with RIPA buffer (50mM HEPES-KOH (pH 7.6), 500mM LiCl, 1mM EDTA, 1% NP40 and 0.7% sodium deoxycholate) and once with Tris-EDTA. Samples were eluted with SDS lysis buffer, RNase A- and proteinase K-treated and crosslinks were reversed at 65 °C overnight. DNA was purified by PCR purification column (Qiagen) and analyzed by qPCR, relative to input. PCR primer sequences are given in Table S5. For ChIP-seq, libraries were generated using the Ultra II library preparation kit (NEB), then sequenced by paired-end sequencing with a 75 cycle high-output Nextseq 500 kit (Illumina).

For reference-normalized ChIP-seq (Orlando et al., 2014), fixed S2 cells were added to fixed SEM cells prior to sonication, at a ratio of 1:4 S2:SEM. After sequencing, reads were adjusted based on the ratio of dm3:hg19 reads in the input and IP samples for each condition.

### ATAC-seq

ATAC-seq was conducted on 5×10^4^ live cells using Nextera Tn5 transposase (Illumina) as previously described (Buenrostro et al., 2013). Libraries were sequenced by paired-end sequencing with a 75 cycle high-output Nextseq 500 kit (Illumina).

### RNA-seq, qRT-PCR and TT-seq

RNA was extracted from 1×10^6^ cells with the RNeasy Mini Kit (Qiagen). For qPCR, reverse transcription was conducted using Superscript III (ThermoFisher Scientific) with random hexamer primers (ThermoFisher Scientific), and cDNA was analyzed by Taqman qPCR, using the housekeeping gene *YWHAZ* for gene expression normalization.

For patient RNA-seq, mRNA was isolated from bulk RNA using the NEBNext Poly(A) mRNA magnetic isolation module (mRNA) and used to generate a strand-specific library using the NEBNext Ultra II Directional RNA Library Prep Kit for Illumina (NEB). Nascent RNA-seq was conducted as previously described (Crump et al., 2021).

TT-seq was conducted as previously described (Schwalb et al., 2016). Briefly, thiouridine-labelled spike-in RNA was generated by in vitro transcription of exogenous plasmid sequences in the presence of 4S-UTP (Jena Bioscience), using the MEGAscript kit (ThermoFisher Scientific). 5×10^7^ SEM cells were treated with 500 µM 4-thiouridine for 5 min, before RNA was isolated by Trizol extraction (ThermoFisher Scientific) with the addition of 60 ng spike-in RNA, then purified and DNase I-treated. Labelled nascent RNA was fragmented briefly by sonication (Covaris), then biotinylated with EZ-link biotin-HPDP (ThermoFisher Scientific) and purified by Streptavidin bead pull-down (Miltenyi). Strand-specific libraries were prepared using the NEBNext Ultra II Directional RNA Library Prep Kit for Illumina (NEB). Libraries were sequenced by paired-end sequencing with a 75 (patient samples) or 150 (nascent RNA-seq) cycle high-output Nextseq 500 kit (Illumina).

### TOPmentation

HT-ChIPmentation was conducted as previously described (Gustafsson et al, 2019). For TOPmentation, protein A-coupled magnetic beads (10 µl) were incubated with 1 µl of the appropriate antibody for 4 hours with rotation at 4 °C in 150 µl Binding Buffer (PBS with 0.5% BSA and 1x protease inhibitor cocktail). Antibodies used are detailed in Table S4. Cells were single-fixed (1% formaldehyde for 10 min) for histone modifications or double-fixed (2 mM disuccinimidyl glutarate for 30 min, then 1% formaldehyde for 30 min) for transcription/chromatin factors. Fixed samples were lysed in 120 µl HT-CM Lysis Buffer (50 mM Tris-HCl pH 8.0, 0.5% SDS, and 10 mM EDTA, 1x protease inhibitor cocktail) and sonicated using a Covaris ME220 (Woburn, MA) to generate 200-300 bp fragments. The sonicated chromatin was then incubated with Triton-×100 at a final concentration of 1% for 10 min at room temperature, to neutralize the SDS in the lysis buffer. The chromatin was pre-cleared for 30 min at 4 °C with rotation using 5 µl protein A-coupled beads to reduce non-specific binding. The antibody incubated beads were washed in 150 µl Binding Wash Buffer (PBS with 0.5% FCS) and the pre-cleared chromatin was then added to the antibody-coated beads before incubation overnight at 4 °C with rotation.

Immunoprecipitated chromatin was washed three times with RIPA buffer (50 mM HEPES-KOH pH 7.6, 500 mM LiCl, 1 mM EDTA, 1% NP-40, and 0.7% Na deoxycholate). Beads were transferred to a second tube between the first and second wash to remove non-immunoprecipitated fragments adhered to the tube. The beads were then washed once with Tris-EDTA and once with 10 mM Tris-HCl pH 8.0. The chromatin was then tagmented by resuspending beads in 29 µl Tagmentation Buffer (10 mM Tris-HCl pH 8.0, 5 mM MgCl_2_, 10% dimethylformamide) and adding 1 µl of transposase (Illumina). Samples were incubated at 37 °C for 10 min and the reaction was terminated by adding 150 µl RIPA buffer. Beads were washed with 10 mM Tris-HCl pH 8.0, to remove any detergent, and resuspended in 22.5 µl ddH_2_O. In order to amplify the tagmented chromatin, 25 µl NEBNext Ultra II Q5 Master Mix (NEB) and indexed amplification primers (125 nM final concentration) were added and libraries were prepared using the following thermal profile: 72 °C 5 min, 95 °C 5 min, (98 °C 10 s, 63 °C 30 s, 72 °C 3 min) x 11 cycles. Library clean-up was performed using Agencourt AMPure XP beads at a 1:1 ratio. Samples were sequenced by paired-end sequencing on a NextSeq 500 (Illumina).

### Next Generation Capture-C

Capture-C was conducted as described previously (Davies et al., 2016; Downes et al., 2022), using 2×10^7^ cells per replicate. Briefly, DpnII-generated 3C libraries were sonicated to a fragment size of 200 bp and Illumina paired-end sequencing adaptors (New England BioLabs, E6040, E7335 and E7500) were added using Herculase II (Agilent) for the final PCR. Indexing was performed in duplicate to maintain library complexity, with libraries pooled after indexing. Enrichment was performed using previously designed Capture-C probes (Godfrey et al., 2019), with two successive rounds of hybridization, streptavidin bead pulldown (Invitrogen, M270), bead washes and PCR amplification using the HyperCapture Target Enrichment Kit (Roche). Samples were sequenced by paired-end sequencing with a 300 cycle high-output Nextseq 500 kit (Illumina). Data analysis was performed using CapCruncher v0.2.0 (https://doi.org/10.5281/zenodo.6326102; (Downes et al., 2022)) and statistical analysis was performed as described (Crump et al., 2021; Godfrey et al., 2019).

### Immunoprecipitation for Mass Spectrometry Analysis

Nuclear extracts from HEK 293 cells expressing a FLAG- and HA-tagged fragment of the MLL protein (amino acids 1101 to 1978; containing the CXXC domain as well as the PHD fingers, referred to as FLAG-C-PHD) were incubated with anti-FLAG M2 beads (Sigma Aldrich) in 50 mM Tris-HCl pH 7.5, 300 mM KCl, and 20% glycerol. Beads were washed in 50 mM Tris-HCl pH 7.5, 200 mM KCl, 0.05% NP-40, and 20% glycerol, and then 3 times with the same again but with 150mM KCl. Bound proteins were eluted with FLAG peptide. Nuclear extracts from normal HEK 293 cells with no expression construct were used as a control. Eluted material was run on a 4-12% NuPAGE gel and stained with colloidal blue. Bands were separately cut out of the FLAG-C-PHD and control lanes and submitted for mass spectrometry.

### Mass Spectrometry Analysis

The protein samples were processed and analyzed at the Mass Spectrometry Facility of the Department of Pathology at the University of Michigan. Each lane was cut into 12 equal slices and in-gel digestion with trypsin followed by LC-MS/MS analysis was performed as described elsewhere (Milne et al., 2010). Spectra were searched using the Comet (version 2022.01 rev. 0) (Eng et al., 2013). Searches were done with the Uniprot Human reference proteome (UP000005640) with reversed sequences appended as decoys (downloaded May 9, 2022) (UniProt, 2021). Each search was done using the following parameters: parent mass error +/- 3 Da; fragment bin ion tolerance +/- 1.0005; tryptic digestion with up to 1 missed cleavage; variable modification of +16 Da on methionine and a fixed modification of +57 Da on Cysteine. Search results were then processed using the Trans-proteomic Pipeline (TPP) tools PeptideProphet and ProteinProphet (Deutsch et al., 2010; Keller et al., 2002; Nesvizhskii et al., 2003). These tools collapse proteins with shared peptide evidence into protein groups.

### Additional Immunoprecipitations and GST pulldowns

Nuclear extracts from HEK 293 cells expressing either FLAG-tagged CXXC or PHD constructs were incubated with anti-FLAG M2 beads (Sigma Aldrich) in 50 mM Tris-HCl pH 7.5, 300 mM KCl, and 20% glycerol. Beads were washed in 50 mM Tris-HCl pH 7.5, 150 mM KCl, 0.05% NP-40, and 20% glycerol. Bound proteins were eluted with FLAG peptide. For GST pull-down assays, 500 ng of GST-fused proteins and 200 ng of purified factors were mixed with glutathione-sepharose 4B beads in binding buffer (20 mM Tris-HCl pH 7.9, 300 mM KCl, no EDTA, 20% glycerol, 0.1% NP-40) and washed in the same buffer. Bound proteins were analyzed by western blotting (for antibodies see Table S4) and/or silver staining.

### TetR cell lines

For the TetR recruitment assay, we used the previously engineered *Tet Operon* (*TetO*) mESC line (Blackledge et al., 2014). The MLL CXXC domain, ENL, PAF1 and DOT1L cDNA sequences were cloned into the original pCAGFS2TetR vector, downstream of the FS2-TetR open reading frame. Plasmids were transfected into mESC using Lipofectamine-2000, and puromycin (1 µg/ml) was used to select for clones stably expressing the TetR construct (confirmed by immunoblot). As a negative control, cells were treated with 1 µg/ml doxycycline for 6 h to disrupt TetR binding at the *TetO* prior to fixation for ChIP.

### Generation of degron cell lines

SEM cell lines were generated with either both endogenous copies of the *PAF1* or *SSRP1* gene fused to *FKBP12^F36V^-P2A-mNeonGreen*, immediately prior to the stop codon. This was achieved by CRISPR/Cas9-mediated homology-directed repair (HDR), as previously described (Hyle et al., 2019). SEM cells were electroporated with two plasmids: pX458, encoding Cas9, an sgRNA sequence targeting the end of *PAF1* or *SSRP1* (see Table S5) and mCherry; and an HDR plasmid containing the *FKBP12^F36V^-P2A-mNeonGreen* sequence flanked by 500 bp sequence with homology to either side of the insertion site. After 24h, mCherry-positive cells were isolated by fluorescence-activated cell sorting to identify cells expressing Cas9, and allowed to grow for 1-2 weeks. mNeonGreen-positive cells were subsequently sorted, to isolate cells that had correctly incorporated the insertion sequence within the open reading frame of *PAF1*/*SSRP1*. After a further 1-2 weeks’ growth, cells with the highest mNeonGreen signal were isolated to enrich for homozygous insertions and plated onto H4100 Methylcult (StemCell Technologies) to isolate clonal populations.

### ChIP-seq/TOPmentation and ATAC sequencing analysis

Following quality checking of FASTQ files by fastQC (https://www.bioinformatics.babraham.ac.uk/projects/fastqc/), reads were trimmed using trim_galore (https://www.bioinformatics.babraham.ac.uk/projects/trim_galore/) to remove contaminating sequencing adapters, poor quality reads and reads shorter than 21 bp. Reads were then aligned to hg19 using bowtie2 (Langmead et al., 2009). Duplicate reads were removed using DeepTools alignmentSieve, with the flag –ignoreDuplicates. For ATAC, the – ATACshift flag was also set to correct for adapter insertion. BigWigs were generated using the DeepTools bamCoverage command, with the flags –extendReads –normalizeUsing RPKM, and visualized in the UCSC genome browser (Kent et al., 2002). Peak calling was performed using MACS2 with the flags -f BAMPE -g hs or LanceOTron (Hentges et al., 2021) where any peaks with a score of less than 0.5 were discarded.

### RNA-seq analysis

Reads were subjected to quality checking by fastQC and trimming using trim_galore to remove contaminating sequencing adapters, poor quality reads and reads shorter than 21 bp. Reads were then aligned to hg19 using HISAT2 (Kim et al., 2019) in paired end mode using default parameters. Gene expression levels were quantified as read counts using the featureCounts function from the Subread package (v2.0.2) (Liao et al., 2013) with default parameters. The read counts were used to identify differential gene expression between conditions using the DESeq2 (v3.12) (Love et al., 2014) package. For TT-seq, spike-in RNA levels were quantified by mapping to a custom genome using featureCounts, and used to normalize the output of DESeq2. For metagene analysis, each transcript was scaled by binning the normalized coverage into 600 bins. Upstream and downstream flanking regions (2 kb) were split into 200 equal length bins. Transcripts were split into quartiles based on length, and median coverage within each bin for each quartile was determined. The median of all replicates for all conditions was determined and log2 transformed after adding a pseudo count of 1.

## Data availability

All high throughput data have been deposited in the Gene Expression Omnibus (GEO) under accession number GSEXXXXXX. Accession numbers for datasets used from previous publications can be found in Tables S1 and S6.

## Acknowledgements

T.A.M., N.T.C., A.L.S, L.G., S.R. and N.J. were funded by Medical Research Council (MRC, UK) Molecular Haematology Unit grant MC_UU_00016/6. N.T.C. was supported by a Kay Kendall Leukaemia Fund Intermediate Fellowship (KKL1443). A.R. was supported by a Bloodwise Clinician Scientist Fellowship (grants: 14041 and 17001), Wellcome Trust Clinical Research Career Development Fellowship (216632/Z/19/Z), Lady Tata Memorial International Fellowship, and EHA-ASH Translational Research Training in Hematology Fellowship. I.R. is supported by the NIHR Oxford BRC, by a Bloodwise Program Grant (13001) and by the MRC Molecular Haematology Unit (MC_UU_12009/14). R.G.R. was funded by a Leukemia and Lymphoma Society Specialized Center of Research Grant (17403-19) and the NIH (CA178765). Primary haematological malignancy samples used in this study were provided by Blood Cancer UK Childhood Leukaemia Cell Bank.

## Author Contributions

N.T.C., A.L.S., T.A.M. conceived the experimental design. N.T.C., A.L.S., L.G., N.J., S.R., J.K., V.B. and T.A.M. carried out experiments. N.T.C., A.L.S., D.F. and V.B. analyzed and curated the data. N.T.C., A.L.S. and T.A.M. interpreted the data and wrote the manuscript. R.G.R., C.D.A., T.A.M., A.R. and I.R. provided funding and supervision. All authors edited the manuscript.

---

**Figure S1.**
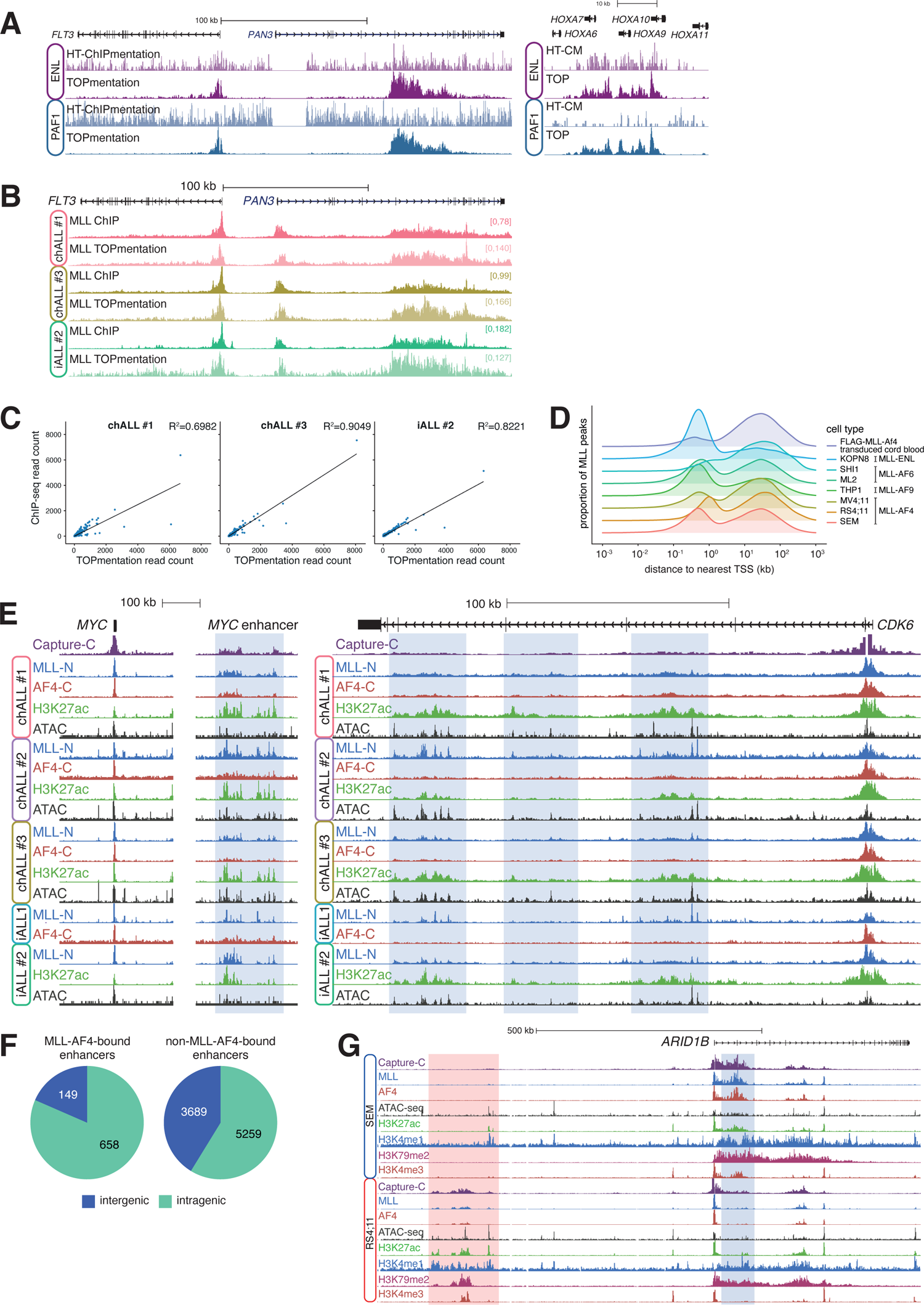
(A) Comparison of ENL and PAF1 signal at the *FLT3* and *HOXA9* loci for high throughput ChIPmentation (HT-CM) (Gustafsson et al., 2019) and TOPmentation (TOP). (B) Comparison of MLL ChIP-seq and TOPmentation at *FLT3* for the indicated MLL-AF4 ALL patients. (C) Correlation of MLL ChIP-seq and TOPmentation signal for the indicated MLL-AF4 ALL patients. (D) Distribution of MLL peaks in the indicated cell lines (or distribution of FLAG tag for FLAG-MLL-Af4 transduced cells), relative to the nearest TSS. Fusion protein characterizing each cell line is indicated. (E) ChIP-seq for MLL, AF4, H3K27ac, and ATAC-seq at *MYC* (left) and *CDK6* (right) in the indicated patient samples. The *MYC* enhancer is approx. 1.7 Mb downstream of the TSS (Crump et al., 2021). Capture-C from the TSS in SEM cells is shown. Common MLL-AF4-bound enhancers are highlighted in blue. (F) Proportion of MLL-AF4-bound and non-MLL-AF4-bound enhancers located within (intragenic) or between (intergenic) genes. (G) ChIP-seq for MLL, AF4, H3K27ac, H3K4me1, H3K79me2 and H3K4me3, and ATAC-seq at *ARID1B* in SEM and RS4;11 cells. Capture-C from the *ARID1B* TSS is shown for SEM and RS4;11 cells. Enhancer elements that show higher activity in RS4;11 and SEM cells are highlighted in red and blue, respectively.

**Figure S2.**
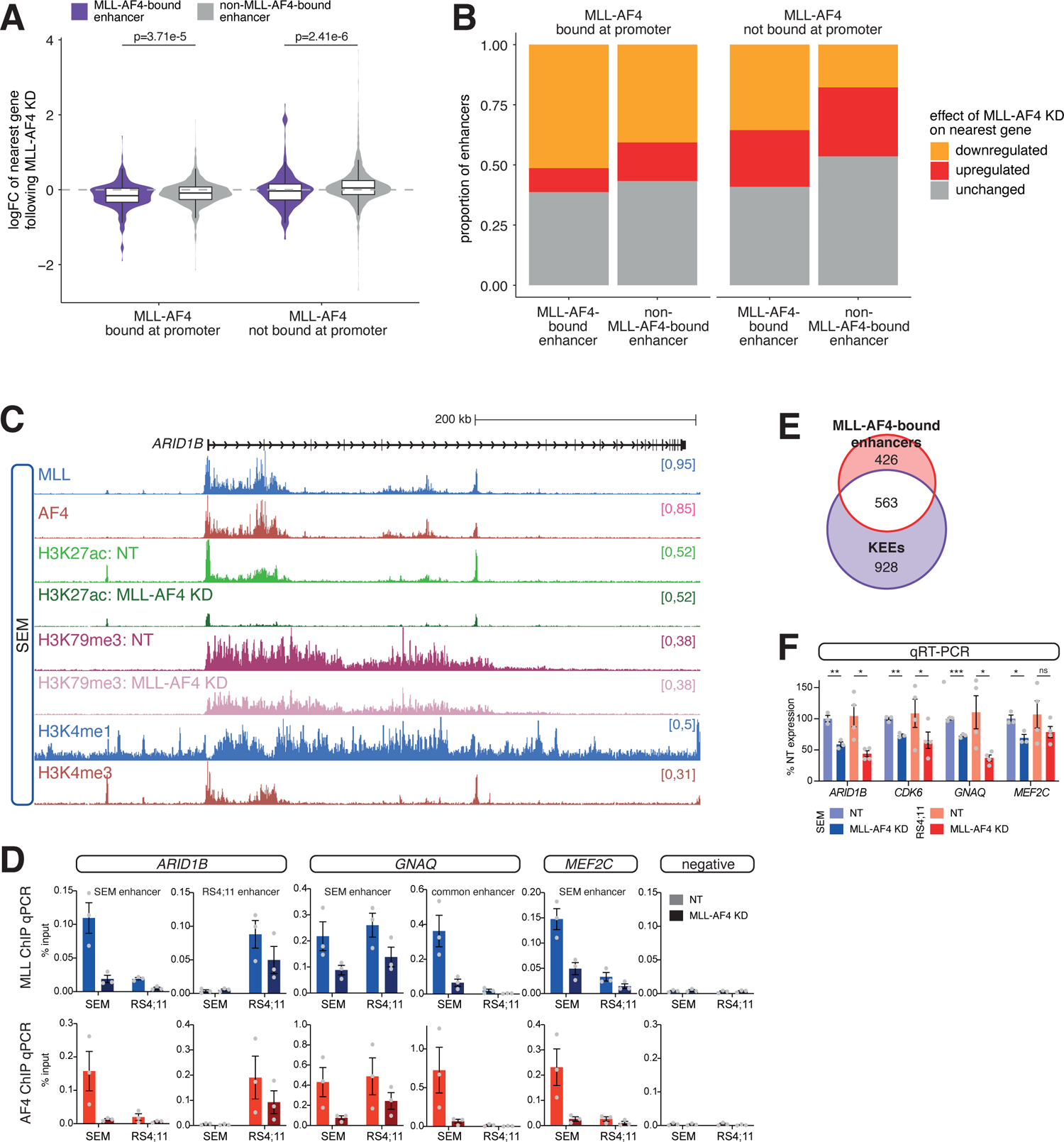
(A) Change in gene expression following KD of MLL-AF4 in SEM cells, separated by the presence or absence of MLL-AF4 within 5 kb of the TSS, for genes associated with an MLL-AF4-bound enhancer and genes associated with an enhancer not bound by MLL-AF4. Statistical significance calculated using a Mann–Whitney U test. (B) Proportion of enhancers associated with genes displaying each transcriptional response to MLL-AF4 KD, separated by the presence or absence of MLL-AF4 within 5 kb of the TSS. (C) Reference-normalized ChIP-seq for H3K27ac and H3K79me3 at *ARID1B* in SEM cells under control (NT) and MLL-AF4 KD conditions. ChIP-seq for MLL, AF4, H3K4me1 and H3K4me3 is shown for context. (D) ChIP-qPCR for MLL and AF4 at the indicated enhancer regions in SEM and RS4;11 cells, under control (NT) and MLL-AF4 KD conditions. Data are represented as mean ± SEM, n=3. (E) Overlap of MLL-AF4-bound enhancers and KEEs in SEM cells. (F) qRT-PCR analysis of gene expression in SEM and RS4;11 cells following MLL-AF4 KD. Data are represented as mean ± SEM, n=3. ns: non-significant; * p<0.05; ** p<0.01; *** p<0.001.

**Figure S3.**
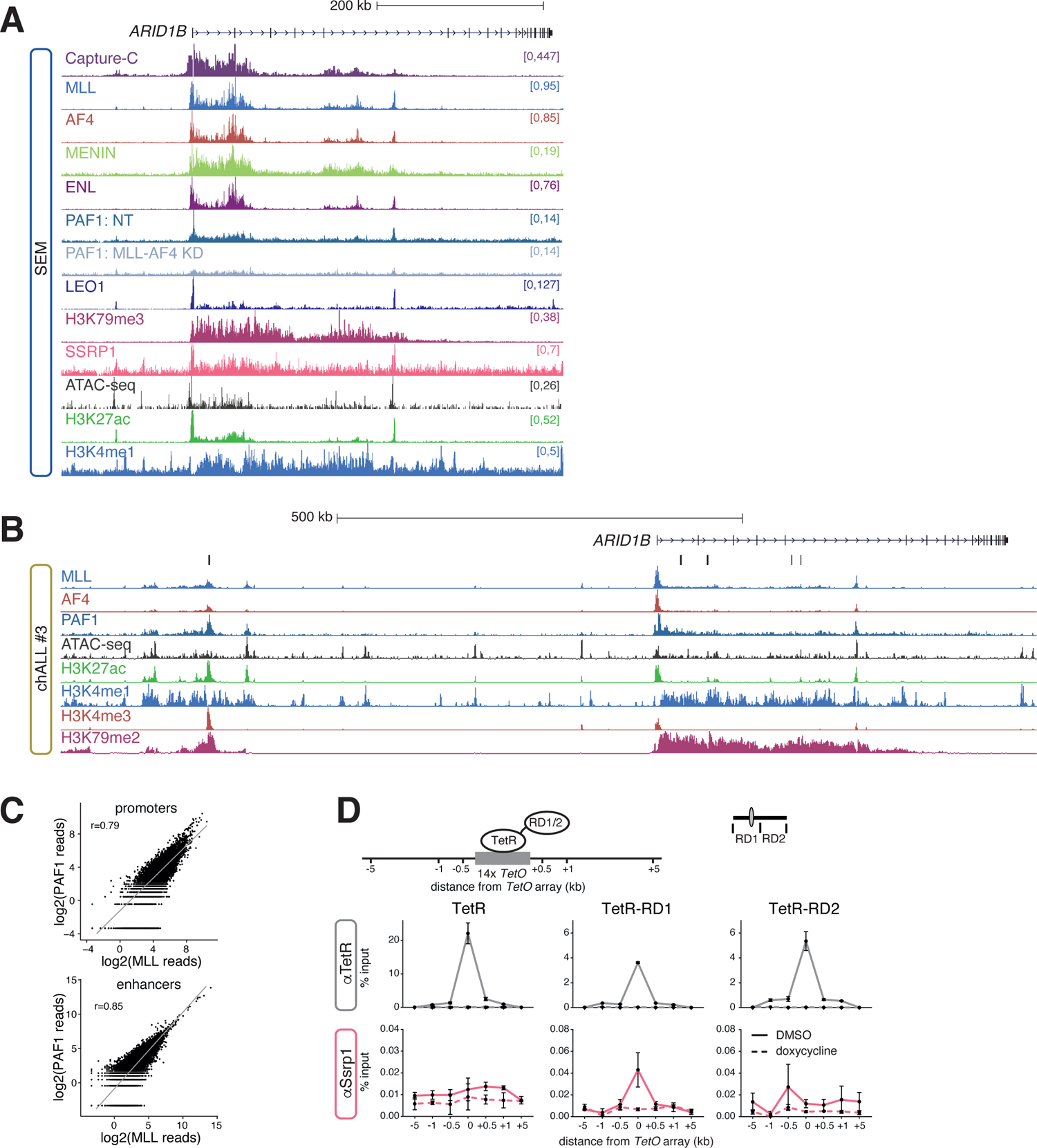
(A) ChIP-seq, ATAC-seq and Capture-C at the *ARID1B* locus in SEM cells. Reference-normalized ChIP-seq for PAF1 in SEM cells under control (NT) and MLL-AF4 KD conditions. The Capture-C viewpoint is the *ARID1B* TSS. (B) TOPmentation and ATAC-seq at the *ARID1B* locus in chALL patient #3. (C) Correlation of MLL and PAF1 TOPmentation at promoters and enhancers in chALL patient #3. (D) ChIP-qPCR for TetR and Ssrp1 at the *TetO* array inserted into mESCs expressing TetR (not fused to another protein), or TetR fused to the RD1 or RD2 fragments of the MLL CXXC domain. Dashed line shows ChIP-qPCR in cells treated with doxycycline for 6h. Data are represented as mean ± SEM, n≥3.

**Figure S4.**
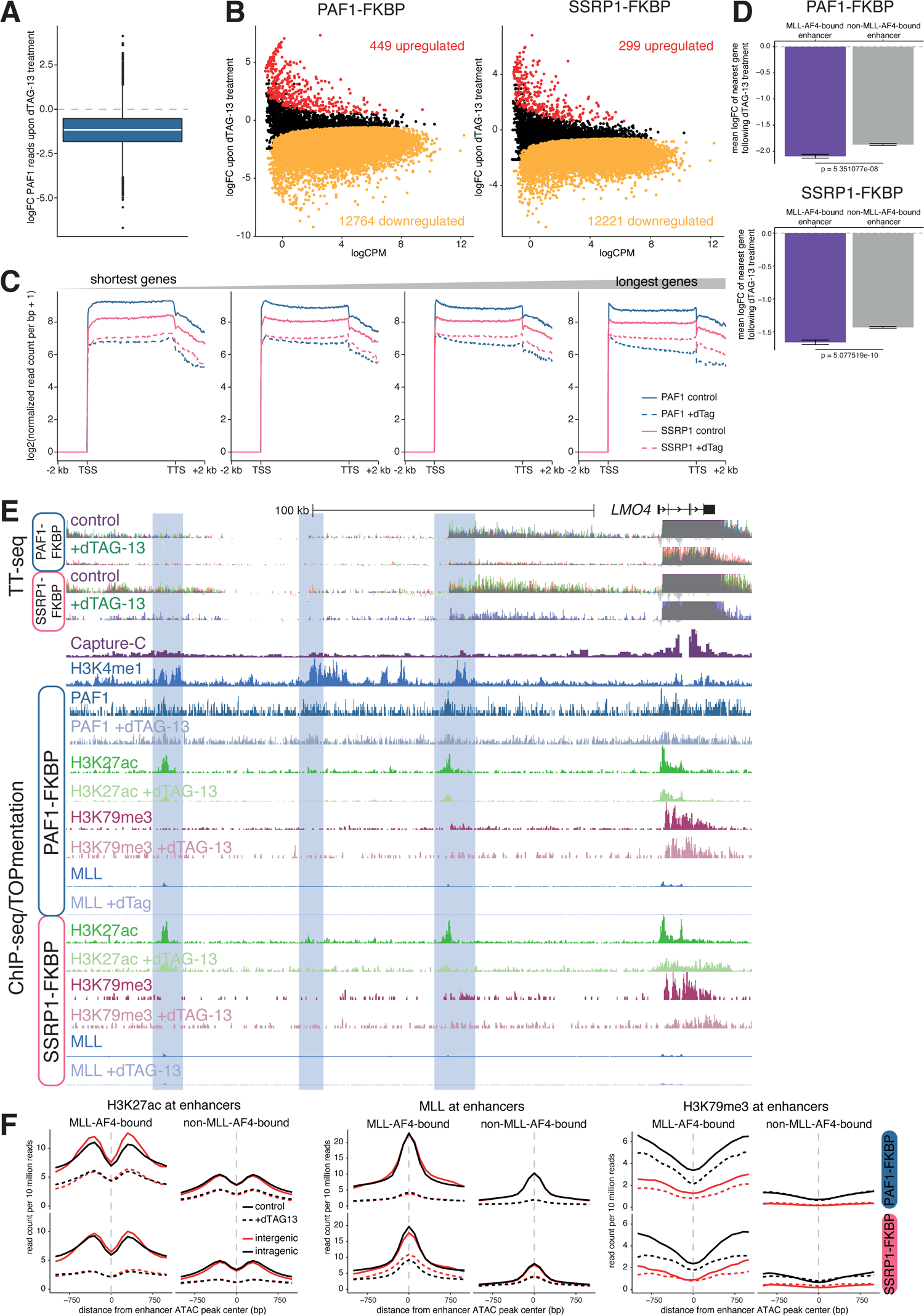
(A) Boxplot of the log2-fold difference in TOPmentation PAF1 reads at PAF1 peaks in untreated or dTAG-13 treated PAF1-FKBP cells. Midline shows median, with upper and lower hinges showing 25th and 75th percentile, respectively. Upper and lower hinges extend to the largest and smallest datapoints within 1.5 times the interquartile range of either hinge. (B) MA plots of TT-seq data showing the effect of dTAG-13 treatment on gene expression in PAF1-FKBP or SSRP1-FKBP cell lines; FDR<0.05, n=3. (C) Metagene profiles of TT-seq levels across gene bodies in PAF1-FKBP or SSRP1-FKBP cell lines, stratified into quartiles by gene length. (D) Mean logFC of expression of genes associated with an MLL-AF4-bound or -unbound enhancer, following dTAG-13 treatment of PAF1-FKBP or SSRP1-FKBP cell lines. (E) TT-seq, reference-normalized ChIP-seq and TOPmentation at the *LMO4* locus in PAF1-FKBP and SSRP1-FKBP SEM cells, with or without the addition of dTAG-13. Capture-C and H3K4me1 ChIP-seq from control SEM cells are shown for reference. The Capture-C viewpoint is the *LMO4* TSS. (F) Mean distribution of H3K27ac, MLL and H3K79me3 at intergenic (red) and intragenic (black) MLL-AF4-bound and non-MLL-AF4-bound enhancers, in PAF1-FKBP or SSRP1-FKBP cell lines under control (untreated) and dTAG13-treated conditions.

**Figure S5.**
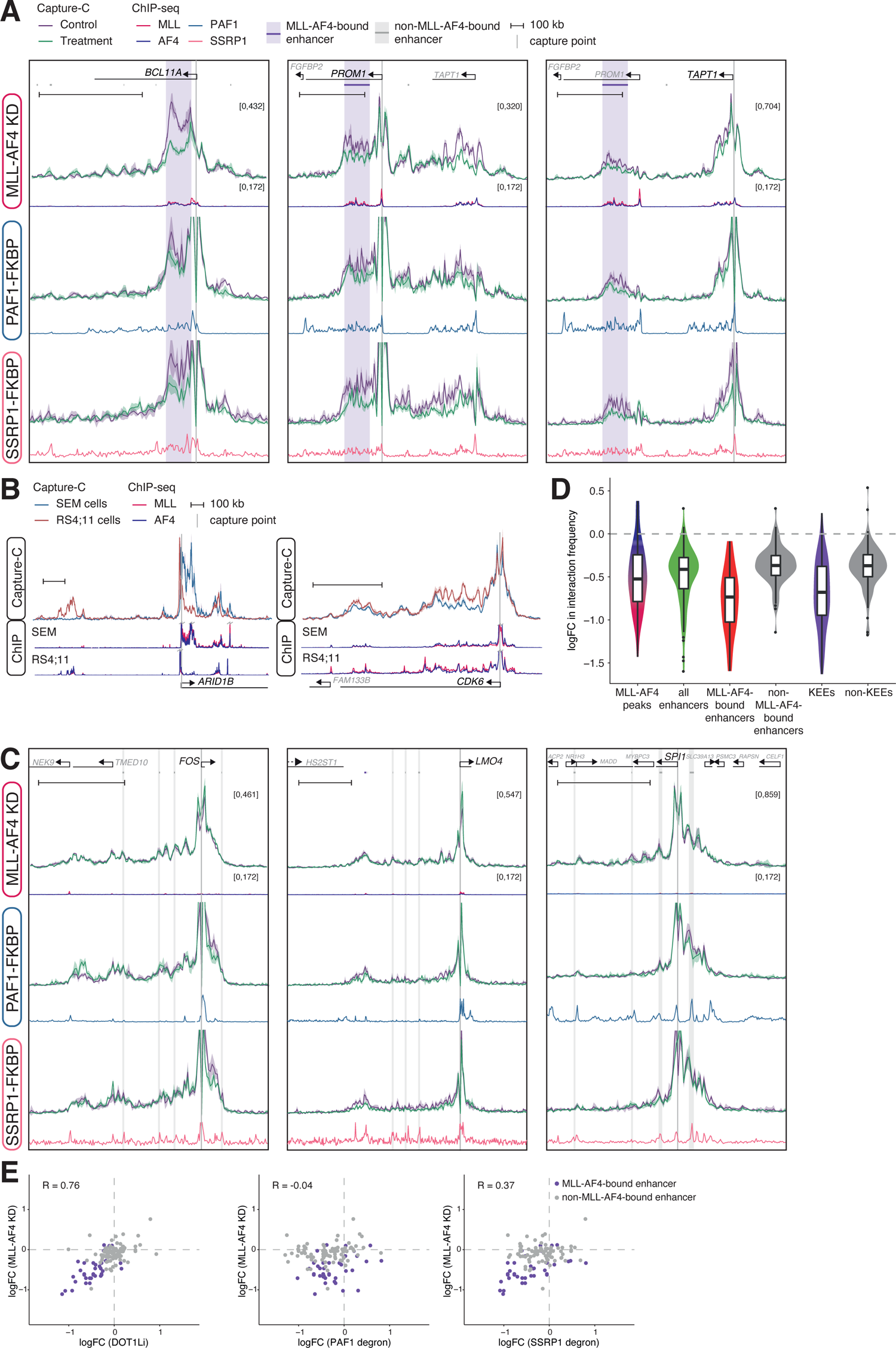
(A) Capture-C from the promoters of *BCL11A*, *PROM1* and *TAPT1* in SEM cells under control (purple) and MLL-AF4 KD (green) conditions (*upper*) or in PAF1-FKBP or SSRP1-FKBP cell lines under control (purple) and dTAG13-treated (green) conditions. Capture-C traces are scaled to emphasize non-proximal interactions. Lines represent mean, shading represents ± SEM, n=3. ChIP-seq traces for MLL, AF4, PAF1 and SSRP1 are shown, along with bioinformatically-annotated MLL-AF4-bound and -unbound enhancers. (B) Overlayed Capture-C from the promoters of *ARID1B* and *CDK6* and ChIP-seq for MLL and AF4, in SEM and RS4;11 cells. ChIP-seq tracks are scaled to emphasize differences in enhancer binding; gray bolts indicate where signal extends beyond the axis. (C) Capture-C from the promoters of *FOS*, *LMO4* and *SPI1* in SEM cells under control (green) and MLL-AF4 KD (purple) conditions (*upper*) or in PAF1-FKBP or SSRP1-FKBP cell lines under control (green) and dTAG13-treated (purple) conditions. Capture-C traces are scaled to emphasize non-proximal interactions. Lines represent mean, shading represents ± SEM, n=3. ChIP-seq traces for MLL, AF4, PAF1 and SSRP1 are shown, along with bioinformatically-assigned MLL-AF4-bound and -unbound enhancers. (D) Change in Capture-C interaction frequency between promoters and the indicated genomic loci, following MLL-AF4 KD. (E) Change in Capture-C interaction frequency between promoters and MLL-AF4-bound (purple) and -unbound (gray) enhancers following the indicated treatments. DOT1Li Capture-C data taken from Godfrey et al 2019.

**Table S1.** High-throughput data collected from MLL-AF4 ALL patient samples.

| Sample | Age at diagnosis | Dataset |  | Source |
| --- | --- | --- | --- | --- |
| chALL #1 | 15 years | ChIP-seq | H3K4me1 | GSE135024 |
|  |  |  | H3K4me3 |  |
|  |  |  | H3K27ac |  |
|  |  |  | H3K79me2 |  |
|  |  |  | MLL | This study |
|  |  |  | AF4 |  |
|  |  | TOPmentation | MLL | This study |
|  |  | ATAC-seq |  | GSE135024 |
|  |  | polyA RNA-seq |  | This study |
| chALL #2 | 6 years | ChIP-seq | MLL | GSE151390 |
|  |  |  | AF4 |  |
|  |  | TOPmentation | H3K4me1 | This study |
|  |  |  | H3K4me3 |  |
|  |  |  | H3K27ac |  |
|  |  |  | H3K79me2 |  |
|  |  | ATAC-seq |  | This study |
|  |  | polyA RNA-seq |  | This study |
| chALL #3 | 3 years | ChIP-seq | MLL | This study |
|  |  |  | AF4 |  |
|  |  | TOPmentation | MLL | This study |
|  |  |  | PAF1 |  |
|  |  |  | RUNX1 |  |
|  |  |  | H3K4me1 |  |
|  |  |  | H3K4me3 |  |
|  |  |  | H3K27ac |  |
|  |  |  | H3K79me2 |  |
|  |  |  | H3K27me3 |  |
|  |  | ATAC-seq |  | This study |
|  |  | polyA RNA-seq |  | This study |
| iALL #1 | 9 months | ChIP-seq | H3K79me3 | GSE135024 |
|  |  |  | MLL | GSE83671 |
|  |  |  | AF4 |  |
| iALL #2 | 12 months | ChIP-seq | MLL | This study |
|  |  | TOPmentation | MLL | This study |
|  |  |  | RUNX1 | This study |
|  |  |  | H3K4me1 |  |
|  |  |  | H3K4me3 |  |
|  |  |  | H3K27ac |  |
|  |  |  | H3K79me2 |  |
|  |  |  | H3K27me3 |  |
|  |  | ATAC-seq |  | This study |
|  |  | polyA RNA-seq |  | This study |

**Table S3.** Gene promoters analyzed by NG Capture-C.

| Gene | Probe coordinates |  |  |
| --- | --- | --- | --- |
|  | Chromosome | Start | End |
| *ARID1B* | 6 | 157099820 | 157100601 |
| *ASH2L* | 8 | 37961940 | 37963209 |
| *BAZ2B* | 2 | 160472165 | 160472912 |
| *BCL11A* | 2 | 60778931 | 60782448 |
| *BCL2* | 18 | 60986641 | 60988918 |
| *CDK6* | 7 | 92462911 | 92463843 |
| *ELL2* | 5 | 95297309 | 95297978 |
| *EP300* | 22 | 41487595 | 41490012 |
| *EZH2* | 7 | 148580112 | 148580317 |
| *FLT3* | 13 | 28674038 | 28675050 |
| *FOS* | 14 | 75744773 | 75746081 |
| *FUT10* | 8 | 33330317 | 33330678 |
| *GADD45A* | 1 | 68150066 | 68151771 |
| *JARID1A* | 12 | 498164 | 499589 |
| *JMJD1C* | 10 | 65226055 | 65226262 |
| *JUN* | 1 | 59249099 | 59250691 |
| *LMO4* | 1 | 87793884 | 87795360 |
| *MBNL1* | 3 | 151986212 | 151986427 |
|  | 3 | 151986424 | 151989271 |
| *MED13* | 17 | 60141972 | 60142862 |
|  | 17 | 60142859 | 60143711 |
| *MEF2C* | 5 | 88180024 | 88180319 |
| *MSL3* | X | 11774388 | 11776889 |
| *MYB* | 6 | 135501733 | 135502770 |
| *MYC* | 8 | 128748253 | 128748439 |
| *OGT* | X | 70752518 | 70753336 |
| *PROM1* | 4 | 16084464 | 16085645 |
|  | 4 | 16077645 | 16078021 |
| *SMYD2* | 1 | 214454064 | 214456235 |
| *SPI1* | 11 | 47399928 | 47400398 |
| *SUPT3H* | 6 | 45345424 | 45345962 |
| *SUZ12* | 17 | 29058515 | 29059304 |
| *TAPT1* | 4 | 16226868 | 16228644 |
| *TET2* | 4 | 106067209 | 106069662 |
|  | 4 | 106066532 | 106067212 |

**Table S4.** Antibodies used in this study.

| **Target** | **Catalogue Number** | **Company** |
| --- | --- | --- |
| H3K4me1 | pAb-194-050 | Diagenode |
| H3K4me3 | 39159 | Active Motif |
| H3K27ac | C15410196 | Diagenode |
| H3K79me2 | 04-835 | Millipore |
| H3K79me3 | C15410068 | Diagenode |
| H3K27me3 | 07-449 | Millipore |
| MLL | A300-086A | Bethyl |
| AF4 | ab31812 | Abcam |
| ENL | A302-268A | Bethyl |
| PAF1 | A300-172A/1 | Bethyl |
| SSRP1 | A303-067A | Bethyl |
| SPT16 | sc-28734 | Santa Cruz |
| HCFC1 | A301-400A | Bethyl |
| CTR9 | A301-395A | Bethyl |

**Table S5.**
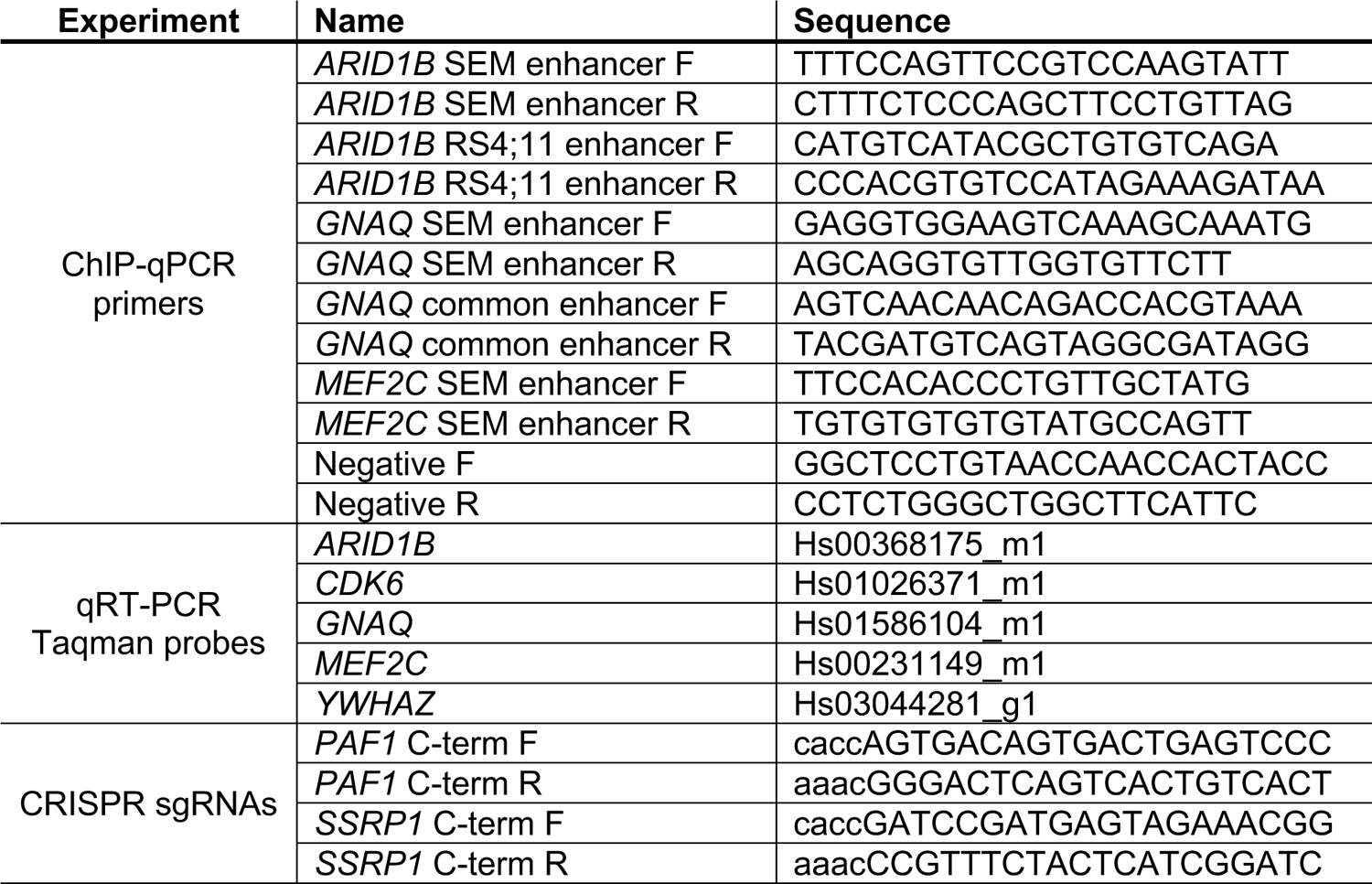
Primers and oligonucleotides used in this study.

**Table S6.**
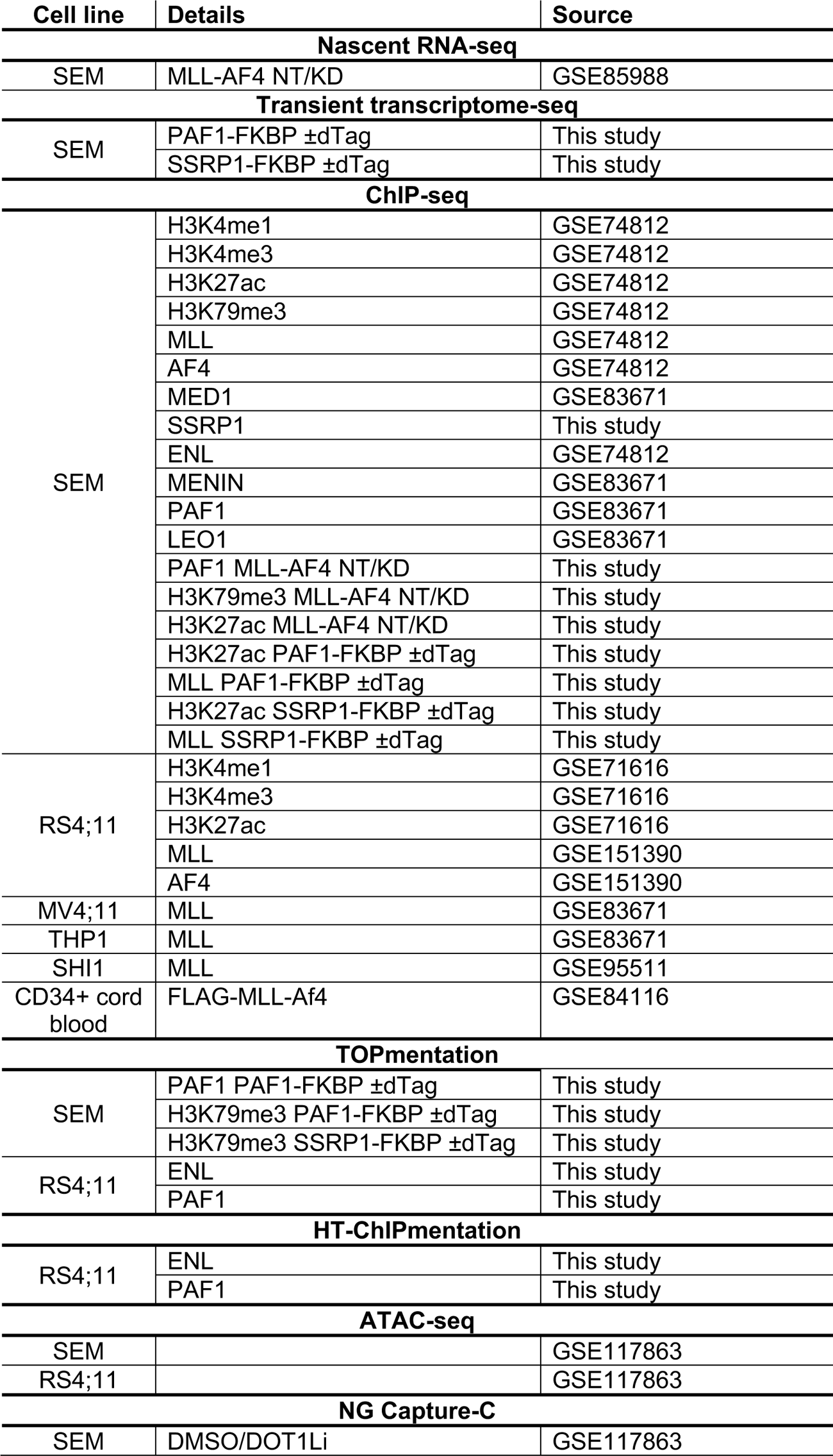
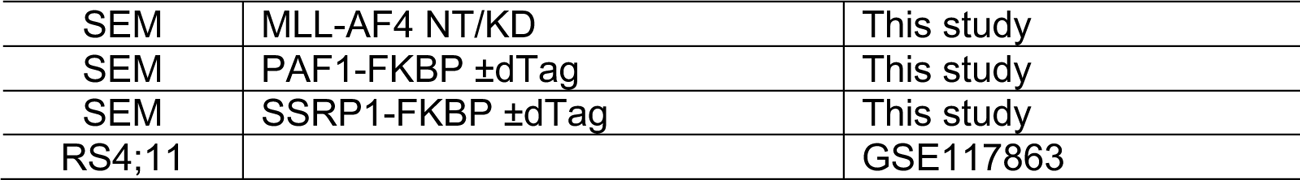
Datasets used in this study.

